# DNA G-quadruplex Profiling Reveals Functional and Mechanistic Role of G-quadruplexes in Skeletal Muscle Stem Cells

**DOI:** 10.1101/2025.02.10.637367

**Authors:** Xiaona Chen, Feng Yang, Suyang Zhang, Xiaofan Guo, Jieyu Zhao, Yulong Qiao, Liangqiang He, Yang Li, Qin Zhou, Michael Tim Yun Ong, Chun Kit Kwok, Hao Sun, Huating Wang

## Abstract

DNA G-quadruplexes (G4s) are non-canonical secondary structures formed in guanine-rich DNA sequences and play important roles in modulating biological processes through a variety of gene regulatory mechanisms. Emerging G4 profiling permits global mapping of endogenous G4 formation. Here in this study, we map the G4 landscapes in adult skeletal muscle stem cells (MuSCs) which are essential for injury induced muscle regeneration. Throughout the myogenic lineage progression of MuSCs from quiescent to activated and further differentiated cells, we uncover dynamic endogenous G4 formation with a pronounced G4 induction when MuSCs become activated and proliferating. We further demonstrate that the G4 induction promotes MuSC activation thus the regeneration process. Mechanistically, we found that promoter associated G4s regulate gene transcription through facilitating chromatin looping. Furthermore, we found that G4 sites are enriched for transcription factor (TF) binding events in activated MuSCs; MAX binds to G4 structures to synergistically facilitate chromatin looping and gene transcription thus promoting MuSC activation and regeneration. The above uncovered global regulatory functions/mechanisms are further dissected on the paradigm of *Ccne1* promoter, demonstrating *Ccne1* is a bona fide G4/MAX regulatory target in activated MuSCs. Altogether, our findings for the first time demonstrate the prevalent and dynamic formation of G4s in adult MuSCs and the mechanistic role of G4s in modulating gene expression and MuSC activation/proliferation.

## Introduction

G-quadruplexes (G4s) are non-canonical secondary structures that can be formed in both single-stranded DNA and RNA sequences enriched with guanines^1,2^. They are typically composed of four tracts of guanines that align in stacked tetra planes stabilized by Hoogsteen hydrogen bonding. Recent studies have revealed that DNA G4s can regulate a wide range of biological processes, e.g., transcription, DNA damage repair and telomere maintenance. Deregulated DNA G4 formation is correlated with many human diseases such as cancer and neurological diseases^3,4^, for example, G4 is found in the promoter of the oncogene *c-MYC* to activate c-Myc expression in lymphoma ^5^. G4s have thus emerged as potential therapeutic targets^3^; various strategies have been developed to target G4 structures, such as small molecule ligands, aptamers, and CRISPR/Cas9 mediated targeting^6–9^. However, currently most of the functional investigations of DNA G4s are limited on individual genes and also in limited cell types due to lack of effective methods to profile endogenous G4 formation in a genome-wide level. Here we profile and investigate the functional roles of endogenous G4s in adult skeletal muscle stem cells.

Skeletal muscle accounts for around 40% of human body mass and adult muscle stem cells (MuSCs, also known as satellite cells) play central roles in muscle tissue homeostasis and regeneration^10–12^. Normally MuSCs remain in quiescence, which is a state of prolonged and reversible cell cycle arrest, in their niche beneath the basal lamina and attached to the myofibers. Upon injury or disease, the MuSCs can go through rapid activation and quickly re-enter the cell cycle, undergoing proliferation as myoblasts. Most myoblasts further undergo differentiation to form new myofibers and fix the damaged muscle, while a subset of cells undergo self-renewal to replenish the pool of quiescent MuSCs. Every stage of the muscle stem lineage progression is tightly controlled by both intrinsic regulatory factors within the cells and extrinsic factors derived from their surrounding niche environment^10,11,13^. Dysregulated MuSCs activities can contribute to the progression of various muscle-associated diseases^10^. Our recent study highlights the regulatory function of RNA G4s in modulating MuSC functions^14,15^. We demonstrate that RNA G4s formed at the mRNA 5’UTR regions inhibit translation and binding of DHX36, a G4 helicase that can unwind the RNA G4 to facilitate mRNA translation during MuSC activation and proliferation. Still, the functional role of DNA G4s and the underlying mechanisms in MuSCs remain unknown.

Earlier studies of DNA G4 were mostly conducted on individual gene locus but recent development of mapping methods such as ChIP-seq and CUT&Tag using a G4-selective antibody BG4 has permitted the global profiling of endogenous G4s and revealed that G4s are prevalently formed at gene promoters in various cell types including cancer cell lines, mouse and human embryonic stem cells and human keratinocyte (NHEK)^16–20^. The successful profiling of endogenous G4 formation has paved the way for the subsequent functional and mechanistic studies. Emerging reports demonstrate that G4s are enriched in open chromatin regions and associated with active transcription, which contrasts with the previously known function of G4s in transcriptional repression^18,19^. Therefore, G4s may contribute to gene regulation through diverse regulatory modes. For example, Lee CY et al. recently demonstrate that G4 formed in the non-template strand increases R-loop formation thus enhances mRNA elongation efficiency and transcriptional yield^21^. Mao S. et al. show that G4 formation sequesters DNA methyltransferase DNMT1 to inhibit promoter methylation and promote transcription^22^. Most interestingly, G4s may modulate chromatin looping between enhancers (E) and promoters (P) to regulate transcription; by acting as structural hubs or recruitment sites for specific trans-acting factors, e.g., CTCF and YY1, G4s may facilitate the formation of chromatin loops, thereby affecting transcriptional output^23–25^. Still, additional factors associated with G4s and other unknown regulatory mechanisms await to be identified; more importantly, the biological significance of G4 regulation remains largely unexplored.

In this study, we for the first time profiled the landscape of endogenous G4 in adult MuSCs undergoing myogenic lineage progression and found the G4 formation is highly dynamic with a pronounced induction when MuSCs become activated. Manipulating global G4 formation by PDS indeed delays MuSC activation/proliferation and muscle regeneration. Moreover, we found that promoter associated G4s globally regulate gene transcription through facilitating DNA looping. Furthermore, we found that G4 sites are enriched for transcription factor (TF) binding events in Activated MuSCs and identified that MAX bound to G4 structures to promote E-P interactions and gene transcription. The above regulatory mechanism was further elucidated on the paradigm of *Ccne1* promoter. Altogether, our results demonstrate the prevalent and dynamic formation of G4s in MuSCs and the positive role of G4s in promoting gene expression and MuSC activation/proliferation through modulating E-P interaction synergistically with MAX.

## Results

### G4 profiling reveals dynamic remodeling of G4s during MuSC lineage progression

To investigate the role of G4 in adult MuSCs, we first profiled the endogenous G4 landscape in MuSCs undergoing lineage progression. From the muscle tissues of 2-month-old PAX7-nGFP mice in which MuSCs were specifically labelled by GFP^26^, the freshly isolated muscle stem cells (FISCs) which are close to quiescent state (Pax7+/MyoD-) were collected by fluorescence-activated cell sorting (FACS); FISCs were further cultured for 24 hours to become activated cells (ASCs-24h) and for 48 hours to be fully activated and proliferating (ASCs-48h, Pax7+/MyoD+); differentiating myoblasts (DSCs, MyHC+) were collected after 72 hours of culturing in growth medium followed by 24 hours in differentiation medium ^14,27^. The above cells were subject to CUT&RUN-seq utilizing the BG4 antibody^28^, to map the global formation of G4s (Fig. 1A, Suppl. Fig. S1A-C). Three biological replicates were conducted and the replicates showed high reproducibility at each stage (Pearson correlation coefficient, r > 0.8) within the same group (Suppl. Fig. S1D). In total we identified 423, 8994, 9421 and 4314 G4 peaks in FISCs, ASCs-24h, ASCs-48h and DSCs, respectively (Fig. 1B, Suppl. Table S1). Motif scanning revealed a high prevalence of G-rich sequences in the G4 peaks identified in ASCs-24h, -48h and DSCs; interestingly, FISC G4s were distinctly enriched for GT repeats which can also form G4 structure as previously reported^29^ (Fig. 1C). Indeed, further analysis uncovered these G4 sequences highly overlapped (69% - 92.7%) with predicted G4 forming sequences (PQS) according to G4 pattern-matching motif scanning (see method, Suppl. Fig. S1E) and enriched PQS signals were observed at G4 formation sites (Suppl. Fig. S1F). Further G4 subtypes analysis showed the identified G4s covered a broad spectrum of G4 structural subtypes, including canonical G4s, long loops, bulges and two quartets^14^ (Suppl. Fig. S1G). Interestingly, the dominant G4 subtype in FISCs was two quartets (35%) which can be formed by the GT repeat sequences followed by bulge subtype (21%), while in ASCs and DSCs, all four subtypes were observed. Altogether, the above results demonstrate the G4 CUT&RUN-seq was successful in capturing the dynamic formation of endogenous G4s in mouse MuSCs undergoing myogenic lineage progression.

**Figure 1.**
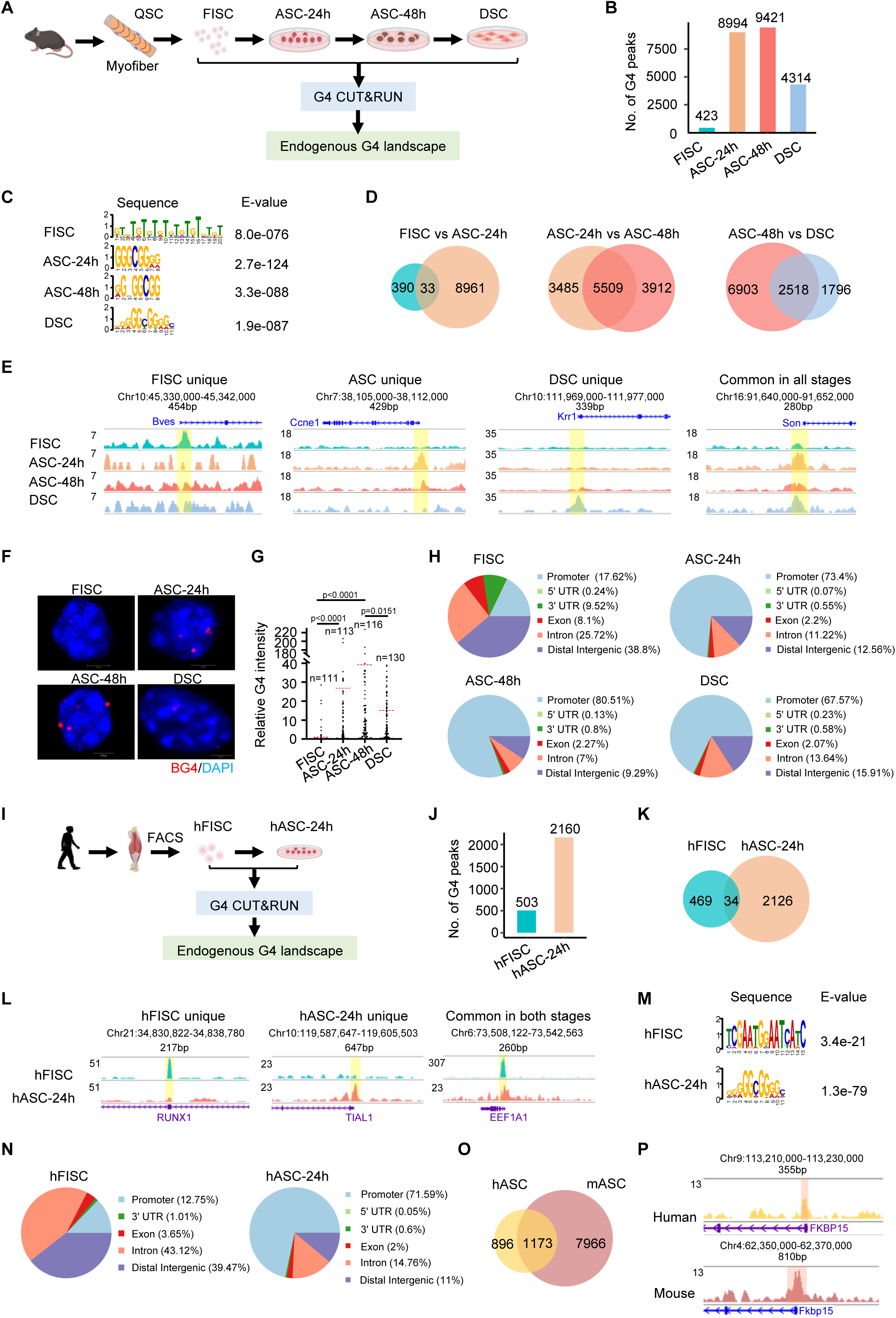
G4 profiling reveals dynamic remodeling of G4s during MuSC lineage progression. (A) Schematic illustration of isolation and collection of adult MuSCs undergoing different stages of lineage progression for G4 CUT&RUN-seq. (B) G4 CUT&RUN-seq was conducted in the above collected cells and the numbers of identified G4 peaks are shown. (C) Motif analysis was performed at the G4 peaks and the top ranked motif sequences are shown. (D) Venn diagrams illustrating remodeling of G4 peaks between two adjacent stages of the MuSC lineage. (E) Genome browser view of the selected stage unique G4 peaks associated with the promoters of *Bves*, *Ccne1*, *Krr1* and *Son* genes. Yellow box highlights regions where G4s are identified. (F) Immunofluorescent staining of DNA G4s was conducted in the above isolated cells after pre-treatment with RNase A and representative images are shown. (G) Quantification of the above BG4 signals per nucleus calculated on the sum projection of the Z-stack. Each point represents a single nucleus, and the red horizontal bars represent the mean from the indicated number of randomly selected nucleus. Kolmogorov-Smirnov test was used to calculate the statistical significance. (H) Genomics distribution of the above identified G4 peaks. (I) Schematic illustration of isolation and collection of human MuSCs for G4 CUT-RUN-seq. (J) The number of identified G4 peaks in the above cells. (K) Venn diagram illustrating remodeling of G4s between human FISC and ASC. (L) Genome browser view of G4 signals across the promoters of *EEF1A1*, *RUNX1* and *TIAL1*genes. Yellow box highlights regions where G4s are identified. (M) The top ranked motifs in the above identified G4s. (N) Genomics distribution of the above identified G4 peaks. (O) Venn diagram comparing the promoter G4s between mASC and hASC. (P) Genome browser view of G4 signal comparison in mASC and hASC across the promoters of *FKBP15* gene. Red box highlights regions where G4s are formed.

To further elucidate the G4 landscape dynamics during MuSC lineage progression, we found a dramatic increase of G4 formation when the cells progressed from FISCs to ASCs (423 vs. 8994) with only 33 shared in the two stages (Fig. 1D-E), suggesting G4 remodeling is associated with MuSC activation. The G4 landscapes in ASCs-24h and ASCs-48h on the other hand were highly similar with 5509 shared peaks. Upon differentiation the G4 number dropped to 4314 in DSCs with 2518 unaltered (Fig. 1D-E). Expectedly, high correlation of G4 signals were observed between ASCs-24h and ASCs-48h (coefficient=0.83) while FISCs and DSCs showed the lowest correlation (coefficient=0.36) (Suppl. Fig. S1H). To substantiate the above findings, BG4 immunofluorescent staining was performed and confirmed the significant increase of the global G4 formation in activated/proliferating MuSCs and the decreased G4 upon differentiation (Fig. 1F-G). Altogether, the above results demonstrate that G4 formation is dynamically associated with the myogenic progression of MuSCs and may play functional roles in the process. Further analysis of the G4 genomic locations uncovered that G4s were mainly formed in the distal intergenic regions (38.8%) and introns (25.72%) in FISCs but primarily in promoters (73.4%, 80.5% and 67.6%) and distal intergenic regions (12.6%, 9.3% and 15.9%) in ASCs-24h, ASCs-48h and DSCs (Fig. 1H), suggesting possible dG4 involvement in transcriptional regulation.

To solidify the G4 induction during MuSC activation, we obtained freshly isolated human MuSCs (hFISCs) collected from human hamstring muscles by FACS (Fig. 1I and Suppl. Fig. S1I-J); the cells were cultured for 24 hours to obtain activated cells (hASCs-24h). G4 CUT&RUN-seq was conducted on the cells and we identified a total of 503 and 2160 G4 peaks in hFISCs and hASCs-24h, respectively, with only 34 overlapped (Fig. 1J-L), indicating a pronounced induction of endogenous G4 formation and a dramatic remodeling when MuSCs became activated, analogous to what was observed in mouse MuSCs. Consistently, the top ranked G4 motifs in hMuSCs were also G-rich sequences (Fig. 1M and Suppl. Fig. S1K-M,); and G4s were mainly formed in distal intergenic regions (39.5%) in hFISCs while primarily in promoters (71.2%) in hASCs (Fig. 1N). Moreover, nearly 60% of the promoter locating G4 peaks in hASCs were shared in mASCs (Fig. 1O-P and Suppl. Table S2), suggesting that G4s may have similar regulatory functions in human and mouse ASCs. Altogether, the above results demonstrate the prevalent and dynamic formation of G4s in both human and mouse MuSCs during myogenic lineage progression and their potential involvement in MuSC fate transitions.

### G4s regulate MuSCs function and adult muscle regeneration

The evident induction of G4 formation in ASCs suggests G4s may promote MuSC activation and proliferation. To test this notion, we treated mASCs with a G4 ligand pyridostatin (PDS) which can bind G4s and sterically hinder the interaction of G4s with other proteins^30–32^ and examined its effect on cell proliferation by EdU assay (Fig. 2A). A significant decrease of cell proliferation upon PDS treatment was observed (Fig. 2B). Similar treatment of PDS on hASCs also resulted in repressed cell proliferation (Fig. 2C), demonstrating that G4 formation promotes MuSC proliferation. To further examine if G4 formation regulates MuSC activity *in vivo* during the injury induced muscle regeneration course, muscles of C57BL/6 mice were injected with BaCl_2_ to induce acute damage and regeneration. MuSCs are known to rapidly activate 1 or 2 days after injury (dpi) and undergo proliferation and further differentiation, with newly formed fibers readily seen by 5 dpi^14^. DMSO (Ctrl) or PDS was injected into the injured tibialis anterior (TA) muscle at 6h and 30h post injury (Fig. 2D). At 5 dpi the muscles were collected and decreased number of Pax7+ (Fig. 2E) and MyoD+ (Fig. 2F) cells were detected on the PDS treated muscles, confirming the impaired MuSC proliferation. Expectedly, the mouse regeneration course was delayed as shown by the reduced eMyHC+ fibers (eMyHC stains the newly formed fibers) at 5 dpi (Fig. 2G); H&E staining at 5 dpi also showed significantly compromised muscle regeneration as the cross-sectional area (CSA) measurement of newly formed fibers indicated a shift to smaller fibers (Fig. 2H-I). Altogether, the above results demonstrate that global G4 formation plays a role in regulating MuCS function and muscle regeneration.

**Figure 2.**
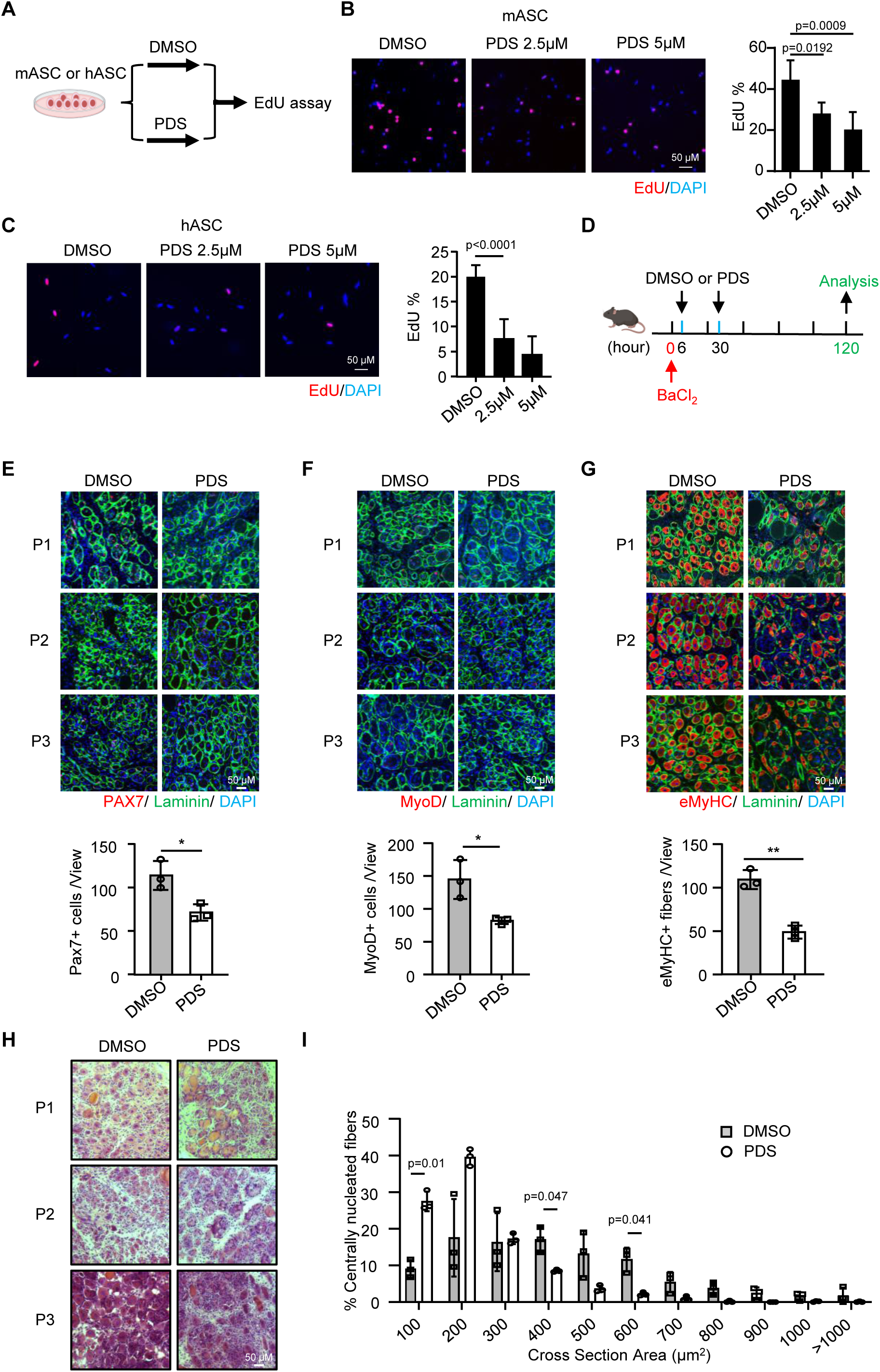
G4s regulate MuSCs function and adult muscle regeneration. (A) Schematic illustration of the PDS treatment with MuSCs. FISCs isolated from mouse or human muscles were cultured for 24h and incubated with PDS (2.5 or 5μM) or DMSO (Ctrl) for another 24h. EdU was then added to the culture medium 4 h before fixation for EdU staining. (B) Representative images of EdU staining on mASCs are shown and the quantification of EdU incorporation percentage was calculated from 10 randomly selected fields. (C) Representative images of EdU staining on hASCs and the quantification are shown. (D) Schematic illustration of the injury-induced muscle regeneration scheme. BaCl_2_ was injected into TA muscles to induce acute injury. PDS or DMSO were injected to the injured muscle at 6h and 30h post injury. The injected TA muscles were harvested at the 5 dpi for the assessment of regeneration process. (E) IF staining of PAX7 (red) and laminin (green) was performed on the TA muscles collected from 3 pairs of mice at 5 dpi. Scale bar=50μm. The positively stained cells were calculated from 10 randomly selected fields in each mouse; n=3 mice per group. (F) IF staining of MyoD (red) and laminin (green) on the above muscles and quantification are shown. (G) IF staining of eMyHC (red) and laminin (green) on the above muscles and quantification are shown. (H) H&E staining of the above muscles. (I) CSAs of newly formed fibers were quantified from the above stained sections and the distribution of fiber sizes is shown, n=3 mice per group. Data represent the average of indicated No. of mice ± s.d.

### Promoter G4 formation regulates gene transcription in ASCs

Next, to study how G4s regulates MuSC activation, we sought to dissect how it modulates gene transcription in ASCs considering the dominant G4 location at gene promoters (Fig. 1H). To this end, RNA-seq was performed in FISCs and ASCs-48h cells; a total of 2787 genes were found up-regulated and 4586 were down-regulated (Suppl. Fig. S2A-D, Suppl. Table S3). By intersecting with the genes with G4 formation induced at their promoters in ASCs (Fig. 3A and Supp. Table S3), we found that indeed these genes displayed differential expressions in ASCs vs FISCs, with 1524 significantly up-regulated (G4-up genes) and 1102 down-regulated (G4-down genes) (Fig. 3B-C), suggesting promoter G4s are correlated with both activated and repressed gene transcription. GO analysis revealed that the G4-up genes were mainly related with cell cycle associated terms, e.g., mitotic cell cycle progression and DNA replication (Fig. 3D), while the G4-down genes were enriched in “negative regulation of cell growth” and “protein modification” etc. (Fig. 3E), indicating that G4 formation modulates the expression of genes related to ASC functions. To further define the targets of G4 regulation, we treated ASCs with DMSO (Ctrl) or PDS and identified the affected genes by RNA-seq (Suppl. Fig. S2E-G). 310 out of the 1524 G4-up genes were significantly down-regulated upon the PDS treatment and defined as the bona fide G4 activated targets (Fig. 3F); GO analysis showed these genes were enriched for “cell cycle phase transition” and “chromatin segregation” etc. (Fig. 3G). *Ccne1* was found to be a prime example of G4-activated targets (Fig. 3H). 69 of the 1102 G4-down genes were up-regulated by the PDS treatment, thus considered as G4 repressed targets (Fig. 3I) and enriched for GO terms such as “negative regulation of muscle contraction” (Fig. 3J). *Etnk2* is shown as a prime target of G4 repression (Fig. 3K).

**Figure 3.**
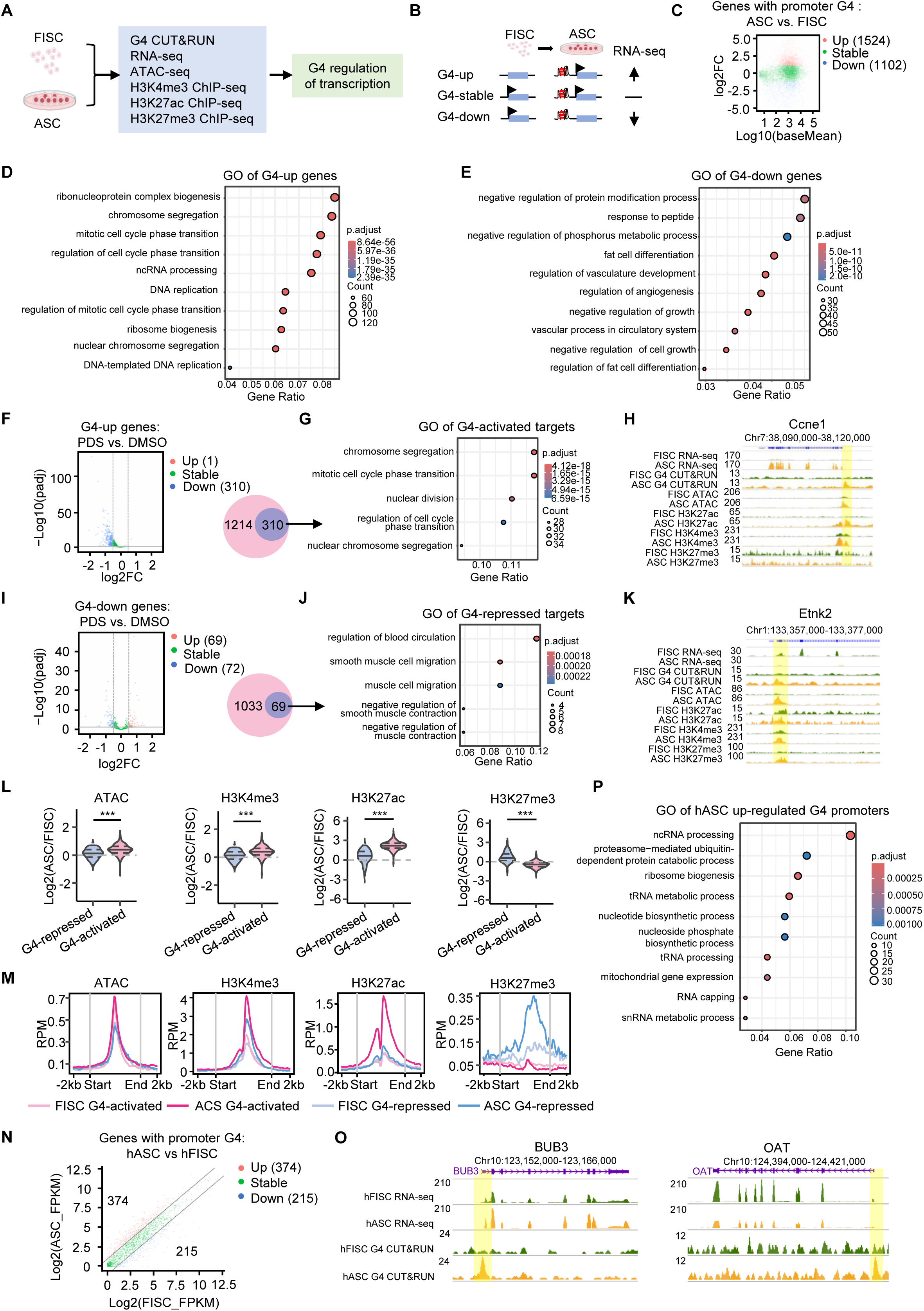
Promoter G4 formation regulates gene transcription in ASCs. (A) Schematic illustration of the analysis integrating multiple datasets for dissecting the global roles of G4s in transcriptional regulation. (B) Schematic illustration of the identification of up- or down-regulated genes by global G4 formation in ASCs vs. FISCs. (C) MA plot showing differentially expressed genes with induced promoter G4 formation in ASCs vs. FISCs. (D) GO analysis of the above G4-up and (E) G4-down genes. (F) ASCs were treated with PDS or DMSO and RNA-seq was performed. Volcano plot showing differentially expressed G4-up genes in ASCs with PDS vs. DMSO treatment. (G) GO analysis of the above defined 310 G4-activated targets. (H) Genome browser view of the G4-activated *Ccne1* locus showing G4 CUT&RUN-seq, RNA-seq, ATAC-seq, H3K27me3, H3K4me3, and H3K27me3 ChIP-seq signals in FISCs and ASCs. Yellow box highlights regions with G4 peaks. (I). Volcano plot showing differentially expressed G4-down genes in ASCs treated with PDS vs. DMSO. (J) GO analysis of the above defined 69 G4-repressed targets. (K) Genome browser view of the G4-repressed *Etnk2* target. (L) Comparisons of the changes of ATAC-seq and H3K4me3, H3K27ac and H3K27me3 ChIP-seq signals at G4-activated and repressed target gene promoters in ASCs vs FISCs. Two-tailed Student’s t-test was used to calculate the statistical significance: ***p < 0.001. (M) Curve plots of the above ATAC-seq, H3K4me3, H3K27ac, H3K27me3 ChIP-seq signals at G4-activated and repressed gene promoters in FISCs and ASCs. For each promoter, the signals are displayed along -2kb to 2kb of transcription start site. (N) RNA-seq was performed in hFISCs and hASCs and intergrated with the G4 CUT&RUN-seq peaks in human MuSCs. Volcano plot showing differentially expressed genes with induced promoter G4 formation in hASCs. (O) Genome browser view of *BUB3* and *OAT* gene loci showing RNA-seq and G4 CUT&RUN-seq signals in hFISC and hASC. Yellow box highlights regions with G4 formation. (P) GO analysis of the above identified G4 up-regulated genes in hASCs.

Next, combining with published ATAC-seq datasets from mouse MuSCs^33^ we found that compared with G4-repressed targets, the G4-activated targets such as *Ccne1* expectedly showed significantly increased chromatin openness at the promoter G4 formation sites in ASCs vs FISCs and this was accompanied with evident increase of H3K4me3 and H3K27ac ChIP-seq signals (Fig. 3H and Fig. 3L-M); upon PDS treatment, the H3K4me3 signals were also significantly decreased (Suppl. Fig. S2H and Supp. Table S4). On the contrary, the G4-repressed targets such as *Etnk2* displayed dramatic increase of the repressive H3K27me3 ChIP-seq signals in ASCs vs FISCs (Fig. 3K and 3L-M). More interestingly, moderate but detectable levels of ATAC-seq and H3K4me3 signals were observed at the G4 formation sites in FISCs prior to the G4 induction in ASCs, in line with the previous report showing chromatin opening occurs before G4 formation^34^. However, H3K27ac and H3K27me3 signals were barely detectable in FISCs and increased dramatically on G4-activated and G4-repressed gene promoters, respectively only upon MuSC activation (Fig. 3M), suggesting the regulatory role of G4s is associated with distinct chromatin modifications. Altogether, the above results demonstrate that global promoter G4 formation promotes MuSC activation/proliferation by regulating targets important for cell cycle progression.

Lastly, we also performed the RNA-seq with human FISCs and ASCs and identified 3603 up-regulated and 2656 down-regulated genes (Suppl. Fig. S2I and Supp. Table S5). Integrating with the G4 CUT&RUN-seq from human MuSC cells, a total of 374 G4-up genes and 215 G4-down genes were identified (Fig. 3N-O), again suggesting G4s function in both promoting and repressing transcription. Moreover, GO analysis revealed that the G4-up genes were mainly enriched for RNA-processing and ribosome biogenesis associated terms (Fig. 3P).

### G4s are enriched at loop anchors and promote loop interactions in ASCs

To further elucidate how G4s activate gene transcription in ASCs, considering their enrichment at promoters and distal intergenic regions (Fig. 1H), we sought to test whether G4s are involved in promoting DNA interactions to facilitate transcription. We thus collected ASC-48h cells and performed Micro-C that enables nucleosome resolution chromosome folding maps^35^ (Fig. 4A). A total of 6,352 loops in 5k resolution was identified by *Hiccups* ^36^ with various subtypes including P-P (promoter-promoter), E-P (enhancer-promoter) and E-E (enhancer-enhancer) loops (Suppl. Fig. S3A and Suppl. Table S6). Integrating with the G4 CUT&RUN-seq data, indeed, we found the G4 signals were enriched at loop anchors (Fig. 4B): 1356 loops possessed G4s at both anchors (dual-G4 loops), 2105 with G4 at one anchor (single-G4 loops) and 2891 without G4 at their anchors (no-G4 loops) (Fig. 4C and Suppl. Table S6). Furthermore, we found dual-G4 loops showed the highest interaction frequency followed by single-G4 loops while the no-G4 loops showed the lowest interaction frequency (Fig. 4D), suggesting that G4 formation at loop anchors may facilitate loop interaction. Consistently, as loop interaction frequency is positively correlated with gene transcription^37^, we found genes with promoters at the dual G4 loop anchors (a total of 2362) showed the highest expression levels compared to those at the single (1701) or no G4 (811) loops (Fig. 4E-G). Altogether, the above findings suggest G4s promote gene transcription through facilitating DNA loop interactions in ASCs. To strengthen our findings, we also took advantage of the Hi-C datasets generated by Wang R et. al. in proliferating myoblast cells^38^. Consistently, we found that G4 signals were enriched at loop anchors and were positively correlated with loop interaction frequency and gene transcription (Suppl. Fig. S3B-H and Suppl. Table S6).

**Figure 4.**
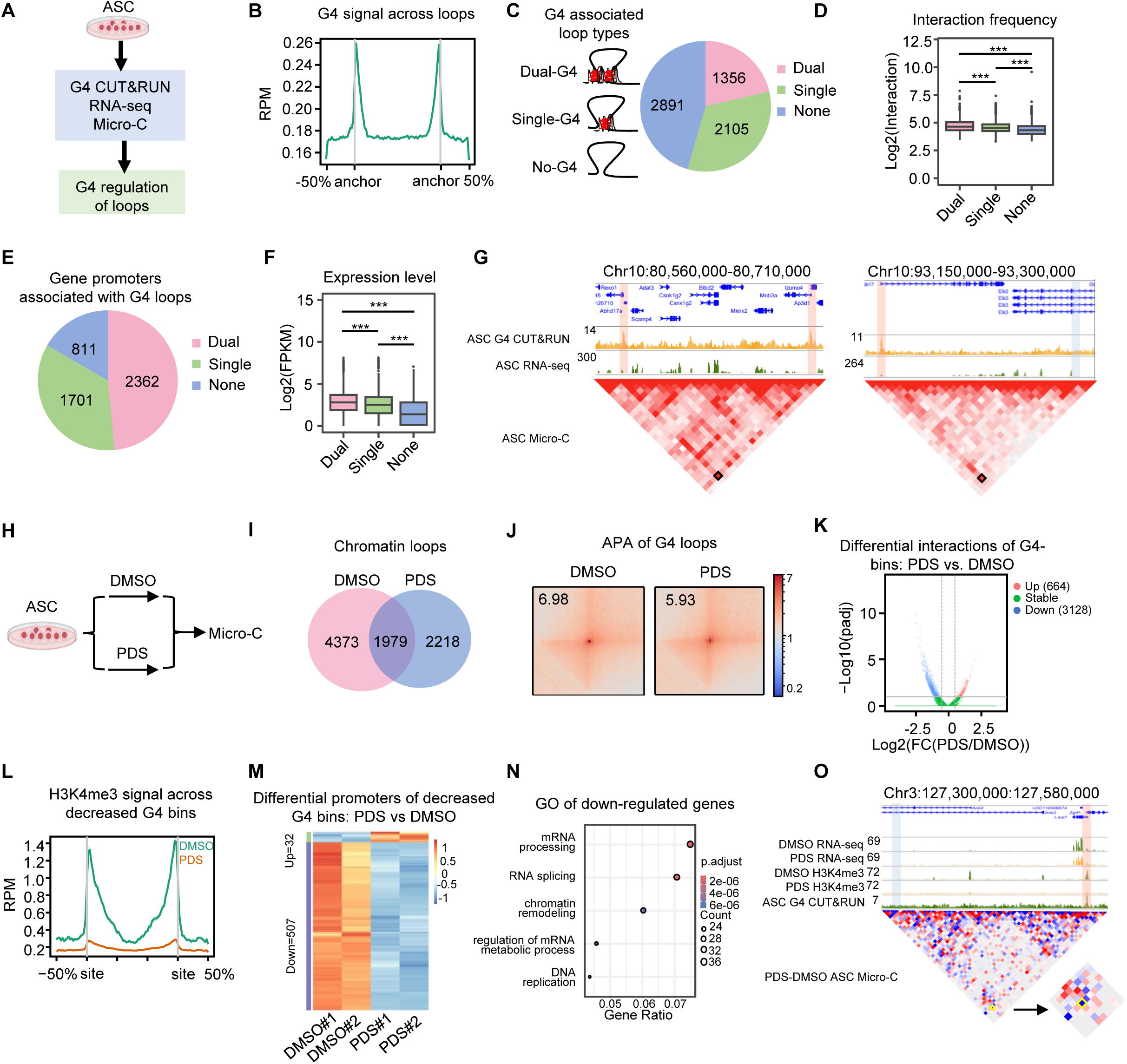
G4s are enriched at loop anchors and promote loop interactions in ASCs. (A) Scheme illustration of integrative analysis to investigate G4 regulation of chromatin looping in ASCs. (B) Chromatin loops were identified by Micro-C and G4 signals across the loops are shown. (C) Schematic illustration of the loop types defined based on the G4 formation at loop anchors and the number of each type in ASCs. (D) Interaction frequency of each type of loop. Student’s t test (two-tailed) was used to calculate the statistical significance: ***P < 0.001. (E) The number of genes promoters located at anchors of each type of loop. (F) Expression level of the above genes in ASCs. Student’s t test (two-tailed) was used to calculate the statistical significance: ***P < 0.001. (G) Genome browser view of G4 CUT&RUN-seq, RNA-seq signals, and Micro-C heatmap of the dual and single loops. The red box highlights the loop anchor with G4, and the blue box highlights the loop anchor without G4. (H) Schematic illustration of PDS or DMSO treatment followed by Micro-C analysis in ASCs. (I) Venn diagram showing loops identified in the above treated cells. (J) Aggregate Peak Analysis (APA) score of the identified G4-containing loops. (K) Chromatin bin interactions with at least one bin region containing G4 were measured, and the up- and down-regulated interactions were identified in ASCs with PDS vs DMSO treatment. (L) H3K4me3 CUT&RUN-seq signals across the above identified decreased interaction regions. (M) Heatmap showing differentially expressed promoters associated with the above decreased interaction regions. (N) GO analysis of the above identified down-regulated genes. (O) Genome browser view of *LARP7* locus showing RNA-seq, H3K4me3 ChIP-seq signals, and Micro-C heatmap comparison in ASCs treated with PDS vs DMSO. The red box highlights the loop anchor with G4, and the blue box highlights the loop anchor without G4.

To substantiate the above findings, we treated ASCs with PDS and assessed the impact of G4 disruption on loop interactions by Micro-C (Fig. 4H). A total of 6352 and 4197 loops were identified in DMSO and PDS treated cells, respectively (Fig. 4I, Supp. Table S6). We found the treatment significantly decreased the interaction strength of the dual and single loops (Fig. 4J and Suppl. Table S6). Consistently, the PDS treatment led to 3128 decreased interactions but a much smaller number (664) of increased interactions, confirming global G4 formation is important for promoting loop interactions (Fig. 4K). To further assess the treatment effect on gene transcription, we examined the gene promoters with decreased loop interactions and found the H3K4me3 signals across the interaction regions were significantly lower upon PDS treatment (Fig. 4L and Suppl. Table S4). Consistently, RNA-seq uncovered 507 down-regulated and only 32 up-regulated genes (Fig. 4M and Suppl. Table S6); and the down-regulated genes were enriched for chromatin remodeling, DNA replication, and mRNA processing etc. (Fig. 4N), in line with the up-regulated genes in ASCs vs FISCs (Fig. 3D). Altogether, the above findings indicate that global G4 formation facilitates DNA loop interactions to promote gene expression thus ASC activation.

### G4-mediated E-P interaction promotes *Ccne1* up-regulation in ASCs

To further illustrate the functional and mechanistic role of G4s in ASC activation/proliferation, we performed in-depth dissection on *Ccne1*, which was identified as a bona fide target of G4 regulation (Fig. 3H). Evident induction of G4 formation was detected at its promoter and H3K27Ac defined enhancers in ASCs accompanied by the formation of E-P interaction and up-regulated gene expression (Fig. 5A). To validate the G4 formation in the *Ccne1* promoter region, the sequence in the promoter was scanned by Quadruplex forming G-Rich Sequences (QGRS) mapper^39^ to identify two G tracts (G4#1 and G4#2) (280nt and 63nt upstream of *Ccne1* transcription start site) with a high G4 formation potential (Fig. 5B). We then performed circular dichroism (CD) spectroscopy with synthetic WT or Mut (several Gs were mutated to As to abolish G4 formation) G4#1 and G4#2 oligos (Fig. 5B); single-stranded DNA oligos were prepared in 150mM KCl or LiCl solutions which were known to promote or repress G4 formation, respectively^15,40^. As expected, both G4#1 and G4#2 WT oligos displayed a strong positive peak signal at 264 nm and also a negative peak signal at 242 nm in K+ but not in Li+ condition (Fig. 5B), which is characteristic of the formation of a parallel G4 structure. On the contrary, Mut oligos displayed weak signals at both wavelengths in K+ and Li+ conditions (Fig. 5B), confirming the absence of G4 structures. To validate the positive regulation of *Ccne1* transcription by the G4 sequences, we cloned the *Ccne1* WT promoter encompassing the WT G4s into the upstream of a luciferase reporter; a mutant reporter was also generated by mutating all the GGG sequences to GTG to disrupt G4 formation (Fig. 5C). Expectedly, significantly decreased luciferase activity was detected in MuSCs transfected with WT vs. Mut reporters (Fig. 5C). Furthermore, the up-regulated expression of *Ccne1* mRNA was confirmed during MuSC activation and proliferation by qRT-PCR (Fig. 5D) and significantly down-regulated upon PDS treatment (Fig. 5E). Altogether, the above findings solidify the positive role of promoter G4s in promoting *Ccne1* transcription during MuSC activation/proliferation.

**Figure 5.**
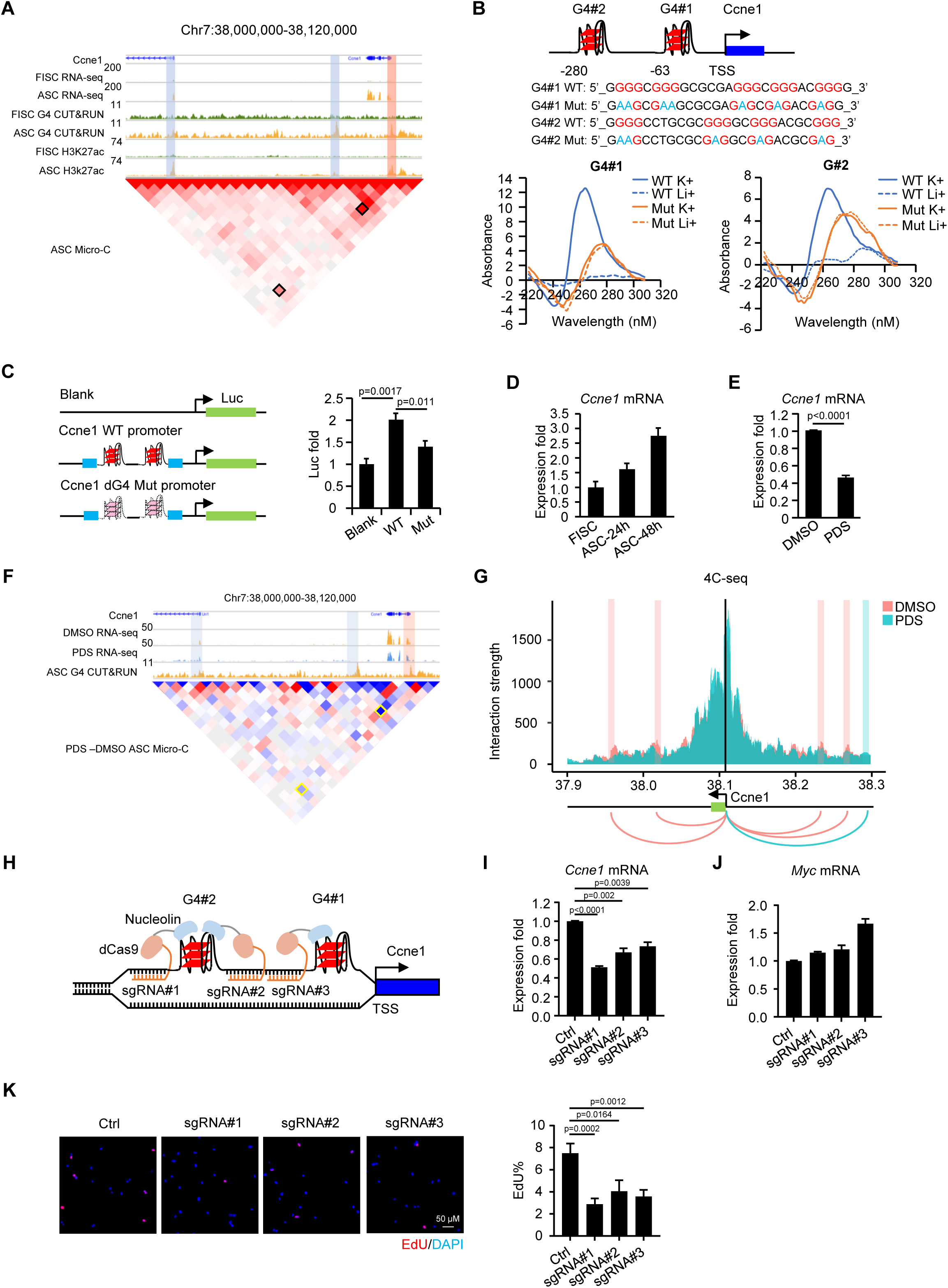
G4-mediated E-P interaction promotes *Ccne1* up-regulation in ASCs. (A) Genome browser view of G4 CUT&RUN-seq, RNA-seq, H3K27ac ChIP-seq signals across *Ccne1* promoter (red bar) -enhancer (blue bar) interaction in FISCs and ASCs and also Micro-C signal in ASCs. (B) Top: sequences of the two WT and Mut G4 sites at the *Ccne1* promoter. All GGGs are highlighted in red. Mutated Gs are highlighted in blue. Bottom: CD spectroscopy was performed using the synthesized WT or Mut DNA oligos under Li+ or K+ conditions. Samples were scanned from 220-310 nm at 25℃. (C) Left: schematic illustration of the dual luciferase reporter. WT or G4 mutated *Ccne1* promoter was cloned into the upstream of firefly luciferase (Luc) ORF. Right: ASCs were transfected with the above generated reporter plasmids or a blank control plasmid and reporter activities were measured (n=3 per group). (D) Expression of *Ccne1* mRNA was examined by qRT-PCR in FISC, ASC-24h or -48h cells. GAPDH was used as the internal control. (E) Expression of *Ccne1* mRNA was examined by qRT-PCR in ASCs treated with DMSO or PDS. GAPDH was used as the internal control. (F) Genomic view of *Ccne1* locus showing RNA-seq, G4 CUT&RUN-seq signals and Micro-C heatmap comparison across the promoter (red box) -enhancer (blue box) interactions in PDS vs DMSO treated cells. (G) 4C-seq was performed in ASCs treated with DMSO or PDS using *Ccne1* promoter G4 as the bait site. The black line represents the viewpoint (VP) at *Ccne1* promoter. Red and green bars represent the interactions between VP and the covered region in DMSO and PDS treatment, respectively. (H) The schematic illustration of tethering nucleolin to *Ccne1* promoter using a CRISPR-dCas9 system. Nucleolin was fused with dCas9 and ASCs were transfected with the dCas9–nucleolin and sgRNAs targeting each of the three *Ccne1* promoter G4s. (I-J) Expression level of *Ccne1* or *MyC* mRNA was examined by qRT-PCR in ASCs infected with lentivirus expressing each of the above sgRNAs or negative control vector. (K) EdU labeling was performed in the ASCs infected with each sgRNA for 4h at 48h post infection and the percentage of EdU+ cells was quantified.

To further elucidate the involvement of E-P interaction in G4 regulated *Ccne1* expression, we found the E-P interaction identified by Micro-C was indeed diminished upon PDS treatment (Fig. 5F). 4C-seq^41^ was also conducted in myoblast cells to examine the alteration of looping at *Ccne1* promoter in high resolution and uncovered the contact frequency of *Ccne1* promoter and enhancer sites was significantly decreased by the PDS treatment (Fig. 5G). Moreover, to pinpoint the role of the identified G4s in regulating *Ccne1* transcription, we employed dCas9/sgRNA to anchor the G4 stabilizing protein nucleolin (NCL)^7^ specifically to G4 loci at the *Ccne1* promoter region in ASCs (Fig. 5H). Three sgRNAs were designed to target both G4#1 and #2 formation sites and transduced into ASCs together with a dCas9-NCL expressing lentivirus (Fig.5H). As expected, *Ccne1* expression was significantly reduced upon NCL tethering to the G4 sites (Fig. 5I), while the expression of other known G4 targets such as MyC was unaffected (Fig. 5J). Moreover, functionally the proliferation of ASCs was also significantly impaired as assessed by EdU assay (Fig. 5K). Altogether, the above findings demonstrate G4s formed at the *Ccne1* promoter region promote *Ccne1* transcription via enhancing the E-P interactions in ASCs.

### MAX interacts with G4s to facilitate E-P interactions and gene transcription in ASCs

Next, to further illuminate how G4s enhance E-P interactions in ASCs, we examined the possibility that G4s may function as a recruiting hub for TFs^32^. By harnessing the rich resources of publicly available TF ChIP-seq datasets generated in C2C12 myoblast cells (Fig. 6A and Suppl. Table S7), our analysis revealed that G4 peaks displayed significantly enriched TF binding events (Fig. 6B). Among these TFs, MAX showed the highest overlapping signals with G4 peaks (49%) followed by YY1 (22%), CTCF (15%), USF1 etc. (Fig. 6B-C); and YY1 and CTCF have been shown to interact with G4 structures in HEK293T and mESC cells^23,24^. MAX is a member of basic helix-loop-helix leucine zipper (bHLHZ) family of TFs and can form homodimers or heterodimers with other family members, including Mad, Mxl1 and Myc^42,43^. To validate the interaction between MAX and G4s, we conducted MAX CUT&RUN-seq in ASCs and found the predominant binding of MAX at promoter regions (66%) and intergenic regions (14%) (Fig. 6D and Suppl. Table S8), similar with the distribution of G4 peaks (Fig. 1H). Interestingly, motif scanning showed that the top ranked MAX binding motif was G rich sequences but not the known canonical MAX binding motif CACGTG (Fig. 6E), suggesting MAX binding is largely G4-dependent in ASCs. Indeed, MAX binding sites were highly colocalized with G4 peaks globally (Fig. 6F) and also enriched at loop anchors (Fig. 6G); moreover, MAX binding and G4 formation displayed a highly positive correlation at loop anchors (Fig. 6H). Altogether, the above results suggest that MAX may be recruited by G4s and function synergistically to facilitate loop interactions. Consistently, we found PDS treatment in ASCs caused a dramatic reduction of MAX binding globally and on loop anchors (Fig. 6I-J), To confirm the direct interaction of MAX with G4 structures, we conducted the microscale thermophoresis (MST) binding assay^44^ using the purified MAX protein and synthesized WT or mutated single-strand G4-forming DNA oligos from the well characterized c-Myc promoter (Fig. 6K). A direct interaction between MAX protein and the WT G4 oligos was detected with a *Kd* of 1.8μM and the binding affinity was significantly impaired when the G4 was mutated with a *Kd* of 21μM (Fig. 6K). To further illuminate the function of MAX in regulating DNA looping in ASCs, we conducted Micro-C in ASCs transfected with siNC or siMAX oligos (Fig. 6L) and found that knocking down MAX altered the chromatin interactions at G4+MAX+ anchor sites; 606 loops showed decreased interactions thus constitute as targets positively regulated by G4/MAX binding; and a smaller number (319) of increased interactions were identified (Fig. 6M). To further assess the effect of MAX knocking down on target gene transcription, we conducted RNA-seq in the above cells and found that 57 were down-regulated and 31 were up-regulated (Fig. 6N), confirming global MAX co-binding is important for promoting G4-mediated loop interactions and gene transcription.

**Figure 6.**
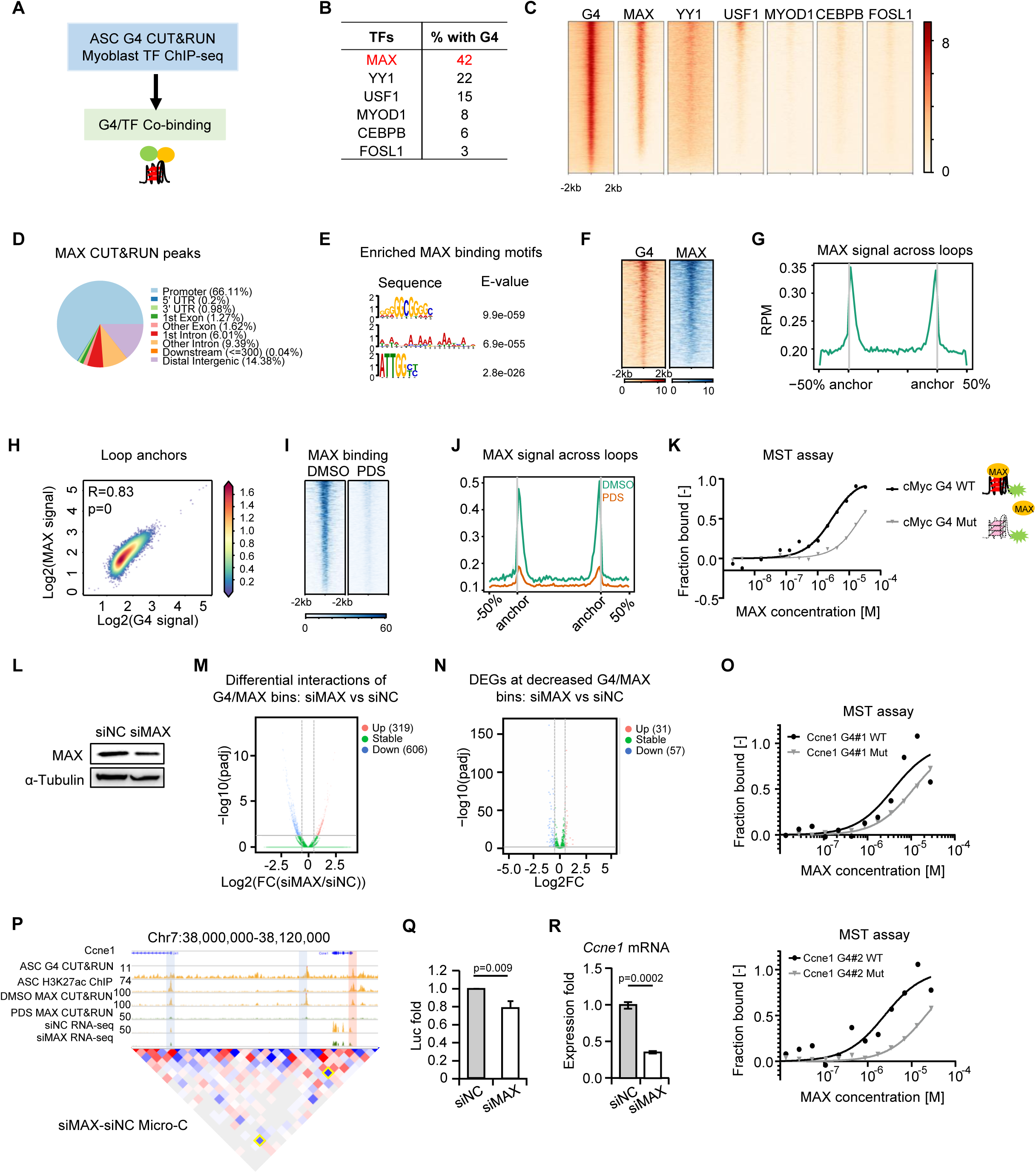
MAX interacts with G4s to facilitate E-P interactions and gene transcription in ASCs. (A) Schematic illustration of the co-binding analysis of G4s and TFs in myoblast cells. (B) Proportion of G4 binding peaks overlapped with the indicated TF binding peaks. (C) Heatmaps of the above TF binding on G4 peaks. (D) MAX CUT&RUN-seq was performed in ASCs and the genomics distribution of MAX binding peaks is shown. (E) The top ranked motifs identified by MEME at the identified MAX peaks. (F) Heatmaps of G4 and MAX binding signals across G4 peaks induced in ASCs. For each peak, the signals are displayed along -2kb to 2kb around the center region. (G) MAX binding signals across loops identified by Micro-C in ASCs. (H) Scatter plot of G4 signals and MAX binding strength at loop anchors identified in ASCs. (I) MAX CUT&RUN-seq was performed in ASCs treated with DMSO or PDS, and heatmaps of MAX binding are shown. (J) MAX signals across loops identified in ASCs treated with DMSO or PDS. (K) MST assay was performed to examine the binding affinities of MAX protein and G4 using purified recombinant human MAX protein and WT or Mut G4 sequences from cMyC promoter. *Kd* was determined for WT and Mut G4s. (L) ASCs were transfected with siMAX or siNC control oligos and MAX knockdown was examined by western blot with α-Tubulin as the normalization control. (M) Chromatin bin interactions with at least one bin region containing G4 and MAX binding were measured and the up- and down-regulated interactions were identified in ASCs transfected with siMAX vs siNC. (N) Differentially expressed genes with promoters associated with the above identified decreased bins. (O) MST assay was performed using MAX protein and WT or Mut G4#1 or G4#2 sequences from *Ccne1* promoter. (P) Genomic view of *Ccne1* locus showing G4 CUT&RUN-seq and H3K27ac signals in ASCs, MAX CUT&RUN-seq signals in ASCs treated with DMSO or PDS, RNA-seq signals in ASCs treated with siNC or siMAX, and Micro-C heatmap comparison across the promoter (red box) and enhancer (blue box) in ASCs transfected with siMAX vs siNC oligos. (Q) ASCs were transfected with the WT *Ccne1* promoter reporter together with siNC or siMAX oligos and the luciferase reporter activities were measured. (R) Expression level of *Ccne1* mRNA was examined by qRT-PCR in ASCs transfected with siNC or siMAX oligos.

To specifically illuminate the MAX regulation on *Ccne1* promoter, we first examined the physical interaction of MAX with the two promoter G4s using MST assay (Fig. 6O). And MAX binding was largely reduced upon PDS treatment at both its promoter and enhancer (Fig. 6P). To further test the regulatory importance of MAX/G4 co-binding in orchestrating *Ccne1* expression, we found that MAX knock-down by siRNA oligos significantly down-regulated the promoter luciferase reporter activity (Fig. 6Q) and *Ccne1* expression (Fig. 6R) in ASCs. Consistently, PDS treatment also decreased the *Ccne1* expression in mASCs (Fig. 5E). Altogether, the above findings identify MAX as G4 binding protein and demonstrate G4/MAX synergistically facilitate loop interactions to promote gene expression in ASCs.

### MAX promotes MuSCs proliferation and adult muscle regeneration

Lastly, to demonstrate the functional role of MAX in ASCs, we found that MAX expression level was expectedly up regulated during MuSC activation and proliferation (Fig. 7A) in accordance with the G4 induction (Fig. 1B). Knockdown of MAX by siRNA oligos significantly impeded ASC proliferation as assessed by EdU assay (Fig. 7B). To further elucidate the functional role of MAX *in vivo* during muscle regeneration, we knocked out MAX in MuSCs using our established *in vivo* CRISPR/Cas9/AAV9-sgRNA genome editing system^45^. Briefly, Adeno-associated virus (AAV) expressing sgRNAs targeting MAX exons were injected into the limb muscles of Pax7^Cas9^ mouse pups at postnatal 7 and 10 days; at 2 months post injection MuSCs were isolated and almost complete deletion of MAX proteins was observed in the isolated ASCs (Fig. 7C-D). The MuSCs were cultured for 2 days for EdU assay; expectedly, a significantly decreased rate of proliferation was observed in the MAX KD vs Ctrl ASCs (11.75% vs 18.4%) (Fig. 7E). To examine the effect MAX KD in muscle regeneration, TA muscles of the 2-month-old mice were injected with BaCl2 and collected at 5 dpi. IF staining of eMyHC showed a significantly decreased number of newly formed fibers in the MAX KD mice (Fig. 7F). A significant delay in muscle regeneration was also observed by H&E staining showing the shifting toward smaller myofibers (Fig. 7G-H). Altogether the above results demonstrate that MAX is essential for MuSC activation/proliferation and adult muscle regeneration, strengthening its functional synergism with global G4 formation.

**Figure 7.**
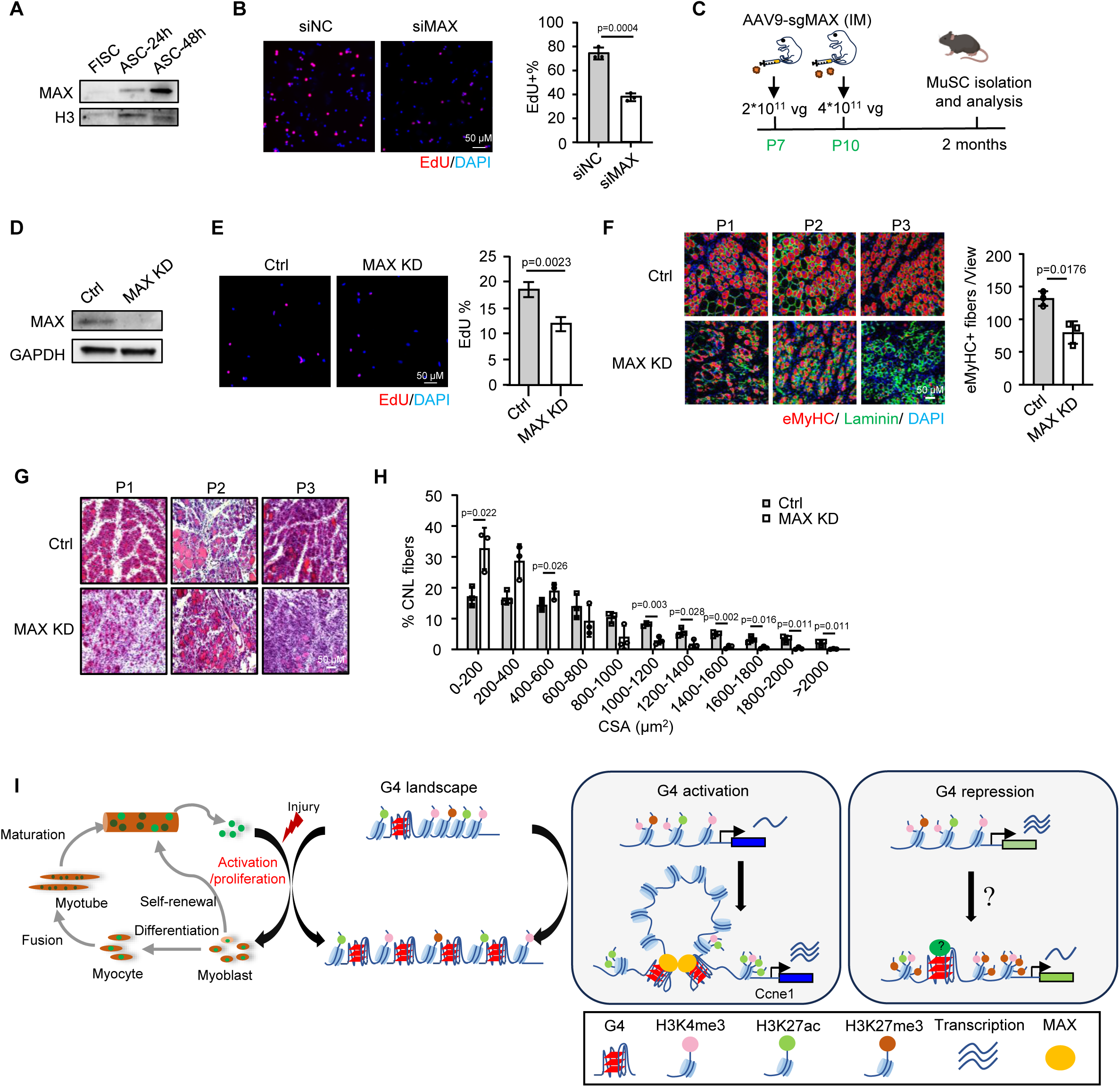
MAX promotes MuSCs proliferation and adult muscle regeneration. (A) MAX protein levels were examined by western blot in FISCs, ASC -24h, ASC-48h with H3 as the normalization control. (B) EdU labeling was performed in ASCs transfected with siNC or siMAX oligos. Representative images of EdU staining are shown and EdU incorporation percentage was calculated. (C) Schematic illustration of the *in vivo* genome editing of MAX in MuSC. AAV9-sgRNA targeting MAX (MAX KD) or negative control virus were injected intramuscularly (IM) to the lower limbs of Pax^Cas9^ mice at postnatal day 7 and day 10. 2 months after the injection MuSCs were isolated and regeneration analysis was also performed on the muscles. (D) Deletion of MAX protein in ASCs isolated from the above generated MAX KD or Ctrl mice was confirmed with GAPDH as the normalization control. (E) ASCs were isolated from the above mice and EdU labeling was performed. Representative images of EdU staining are shown and EdU incorporation percentage was calculated. (F) IF staining of eMyHC (red) and laminin (green) was performed on the TA muscles collected at 5 dpi from the above mice. Nuclei are visualized by DAPI staining (blue). Scale bar=50μm. The positively stained cells were calculated from 10 randomly selected fields in each mouse; n=3 mice per group. (G-H) H&E staining was performed on the above muscles; CSAs of newly formed fibers were quantified and the distribution of fiber size is shown, n=3 mice per group. Data represent the average of indicated No. of mice ± s.d. (I) Schematic illustration of the dynamic remodeling of endogenous G4s in adult MuSC lineage progression and also the functional and mechanistic roles of G4s in MuSC activation/proliferation during muscle regeneration.

## Discussion

In this study, we mapped the endogenous G4 landscape in adult MuSCs undergoing myogenic lineage progression. Our G4 CUT&RUN-seq results reveal that global G4 formation is highly dynamic during this process, with a dramatic increase during cell activation and a significant decrease during myoblast differentiation. Manipulation of G4s using PDS demonstrates that global G4 formation promotes MuSC activation/proliferation to enhance mouse muscle regeneration. Integrative multiomic analysis further reveals that G4s are enriched at gene promoters and positively regulate the transcription of cell cycle and proliferation related genes. Furthermore, we found that G4 enhances promoter transcription through recruiting TFs such as MAX to modulate enhancer-promoter looping interactions. The above described G4 regulatory function and mechanism were dissected in-depth on the *Ccne1* locus. Altogether, our findings have for the first time uncovered the dynamic remodeling and functional roles of endogenous G4s in MuSCs and identified previously unknown mechanisms of G4-mediated transcriptional regulation (Fig. 7I).

Profiling the formation of endogenous G4 formation in cells is the prerequisite for studying G4 biological functions but this has not been possible until recent emergence of G4 genome profiling using BG4 antibody. Our G4 CUT&RUN is proven to be successful in profiling global G4 formation in MuSCs. The results uncovered very limited G4 formation in FISCs when MuSCs are close to quiescence. Upon activation, when cells re-enter the cell cycle, G4 formation is drastically induced and remains high when cells proliferate; subsequently, when cells exit the cell cycle to differentiate, G4 formation decreases. These dynamic changes suggest a correlation between G4 formation and MuSC activation/proliferation. Although it is not our focus to investigate how G4 formation is regulated, various mechanisms have been suggested ^1^. For example, active transcription can facilitate the dissolving of DNA duplex and G4 formation ^34^; other biological processes that facilitate the generation of single-stranded DNA such as DNA replication may also correlate with the G4 formation^28^. Moreover, G4 interacting proteins/helicases that can regulate G4 folding/unfolding may also actively regulate G4 formation^1^.

The dramatic induction of G4 formation during MuSC transition from quiescence to activation/proliferation triggered us to investigate the functional roles of G4s in this process. As expected, increased G4 formation appears to promote MuSC activation/proliferation as functional manipulation of G4 structures using PDS influences cell proliferation both *in vitro* and *in vivo* during mouse muscle regeneration. This is in line with existing reports showing G4 manipulation with various G4 ligands can effectively inhibit cancer cell proliferation and tumor growth, suggesting a general function of global G4s in promoting cell proliferation. Nevertheless, G4s may play different roles in other stages of the myogenic lineage progression. For example, the decreased G4 formation during myoblast differentiation indicates global G4s may function negatively in this fate transition; consistently, a recent study profiling the G4 formation in chicken embryo-derived myoblast cells demonstrates decreased G4 formation during myogenic differentiation^20^.

Mechanistically, we dissected how G4s regulate gene transcription both globally and on the *Ccne1* paradigm locus. Early studies investigating the gene regulatory role of G4s primarily used *in vitro* methods such as ectopic reporter assays on individual loci and the results depicted DNA G4 structures as roadblocks to transcription machinery, thereby suppressing gene expression. However, recent advancements following genome-wide mapping of endogenous G4s have demonstrated that G4s are frequently enriched at promoters of actively transcribed genes, where they act as transcriptional enhancers rather than suppressors^16,19,32^. Several studies suggest G4s serve as non-canonical docking sites for TFs to facilitate RNA polymerase recruitment and increase transcription^19,32^; for example, inserting a G4-forming sequence from the KRAS promoter into the MYC promoter using CRISPR-Cas9 mediated genome editing leads to increased MYC expression^46^. Moreover, Marshall P.R. et al showed that G4s can both silence and activate gene transcription in neuron cells^47^. Therefore, promoter G4s may play pleiotropic roles in gene transcription depending on cell contexts and biological processes. Our integrated multiomic data analyses suggest that in ASCs promoter G4s are correlated with both activated and repressed gene expression, which is also associated with different histone modifications. Of note, we observed ATAC-seq and H3K4me3 signals in FISCs prior to G4 induction in ASCs, demonstrating chromatin openness may be a prerequisite of G4 formation. This is consistent with previous finding that G4 formation is coupled to the establishment of accessible chromatin and does not require active transcription^34^. Moreover, in FISCs, moderate but clear H3K4me3 signals were observed at the promoters of both G4-activated and G4-repressed genes prior to G4 formation, while H3K27ac and H3K27me3 signals were very weak. Upon G4 formation in ASCs, it appears that H3K27ac or H3K27me3 modifications determine whether the G4 acts as a transcriptional activator or repressor. Consistently, recent studies also show the enrichment of H3K4me3 and H3K27ac signals at G4 sites is associated with active transcription in cancer cells and lung fibroblast cells^48,49^. Our findings thus highlight the importance of epigenetic context in determining the functional role of G4s, and the epigenetic cofactors associating with G4s can be further investigated in the future.

Furthermore, our findings uncover the mechanistic roles of G4 structures in modulating 3D genome organization in particularly chromatin looping. Despite several emerging reports showing G4 involvement in 3D regulation, the underlying mechanisms remain incompletely understood. We showed that G4s are enriched specifically at chromatin loop anchors to promote loop formation; of note, G4 enrichment appears to be higher at P-P loops compared with E-E and E-P loops in ASCs (Suppl. Fig. S3A). Further dissection identified MAX as a previously unknown TF that is recruited to G4 binding sites to mediate G4 regulation of looping. Direct physical binding was detected between MAX protein and G4 structures. Moreover, we showed that both G4 and MAX are enriched at chromatin loop anchors to synergistically regulate loop formation. MAX binding appears to be largely dependent on G4s since the top ranked MAX binding motifs are G-rich sequences but not its canonical binding motif. Furthermore, MAX binding is also enriched at promoter regions and may primarily regulate P-P looping together with G4, which is different from CTCF that is highly enriched at the intergenic enhancer region to mediate G4 regulation of E-P looping. Disrupting the G4/MAX interactions with PDS or MAX knockdown led to decreased chromatin interactions, providing functional evidence that G4/MAX acts synergistically in modulating DNA looping. Phenotypically, we also demonstrated that knockdown of MAX impaired ASC proliferation and reduced mouse muscle regeneration both *in vitro* and *in vivo*, consistent with the repressive effects of PDS treatment. Therefore, functionally and mechanistically we demonstrate the synergism of G4 and MAX, highlighting a novel mechanism underlying G4 functions in cells. To further elucidate our theory, in-depth dissection was carried out on *Ccne1* locus and our results demonstrate it is a bona fide target of G4/MAX regulation/function in ASCs. Both PDS treatment and site specific G4 manipulation with CRISPR-dCas9 mediated tethering of nucleolin resulted in impaired DNA looping and gene transcription. Site specific G4 manipulation is emerging as potential therapeutics in diseases, which aims to target individual G4 structure with low off-target risks and unwanted side effects. In addition to CRISPR-dCas9 mediated G4-stabilizing compounds, other strategies including ligand-antisense oligo nucleotides (ASO) conjugate^50^, aptamer-ASO conjugate^51^ have been developed recently to increase the specificity of G4 targeting. It will be interesting to test if site specific G4 targeting can be used to enhance muscle regeneration.

## Methods

### Mice

All animal handling procedures and experiment ethics approval were granted by the CUHK AEEC (Animal Experimentation Ethics Committee) under the Ref No. 23-288-GRF, No. 24-150-MIS, No. 23-287-BRD and No. 24-156-BRD. The mice were maintained in the animal room with 12h light/12h dark cycles, at a temperature of 22-24°C, and a humidity level of 40-60% within the animal facility in CUHK. The Tg: *Pax7-nGFP* mouse strain^26^, was kindly provided by Dr. Shahragim Tajbakhsh. The C57BL/6 wildtype mice were acquired from LASEC (Laboratory Animal Services Centre) at CUHK. The Pax7^Cas9^ mouse was generated by crossing homozygous Pax7^Cre^ mice with the Cre-dependent Rosa26^Cas9-EGFP^ knockin mice (B6;129-Gt (ROSA)26Sortm1(CAG-cas9*-EGFP)Fezh/J). The heterozygous offspring were utilized for *in vivo* genome editing experiments as previously described^45^.

### Animal procedures

For BaCl_2_ induced muscle regeneration, the tibialis anterior (TA) muscles were injected with 50 µl of 1.2% BaCl2 (dissolved in sterile demineralized water)^14^. At the indicated time points, mice were euthanized, and muscles were snap frozen for section. For the PDS treatment post induced acute muscle injury, 50 µl of 1.2% BaCl2 (dissolved in sterile demineralized water) was injected into TA muscle of age-appropriate mice. 6 and 30 hours after the injury, 50 µl of 5nmol PDS (dissolved in DMSO) or DMSO (negative control) was injected into the injured TA muscle. Muscles were harvested at designated time points for subsequent histological or biochemical studies. For the AAV injection, the Pax7^Cre^ x Cas9 mice received intramuscular injections of the AAV-sgRNA at 2 different time points: 2 x 10^11^ vg AAV per leg at postnatal day 7 (P7) and 4 x 10^11^ vg AAV per leg at P10. Mice were subsequently analyzed at 8 weeks post injection.

### Human muscle tissue

Hamstring muscle samples were collected during orthopaedic surgery with informed consent from young female patients in the Hong Kong cohort. The informed consent was obtained from the legally acceptable representative. Ethical approval was granted by the Joint Chinese University of Hong Kong-New Territories East Cluster Clinical Research Ethics Committee (Ref No. 2021.255-T). Exclusion criteria were myopathy, hemiplegia or hemiparesis, rheumatoid arthritis or other autoimmune connective tissue disorders, cancer, coronary heart disease, inability to consent, or major surgery in the previous 3 months.

### Satellite cell sorting

Skeletal muscle tissue from mouse hindlimb and forelimb muscles were dissected, gently minced with blades and digested with Collagenase II (1000U/ml) in Ham’s F10 for 90 minutes in a shaking water bath at 37°C. Digested tissue was then washed twice with rinsing media (10% HS, in Ham’s F-10) and centrifuged at 700 xg at 4°C for 5 minutes. Second digestion was performed by adding Collagenase II (1000U/ml) and Dispase (11U/ml) in rinsing media and incubated in shaking water bath at 37°C for 30 minutes. Digested tissue was passed through a 20-gauge needle for 12 times and filtered through a 40 μm filter followed by spinning at 700 xg for 5 min at 4°C. For mouse muscles, MuSCs were then sorted out by FACS Aria Fusion (BD) and collected as GFP+ groups. For human muscles, cells were stained with antibodies and FITC-(CD45-, CD31-, CD34-) APC+(CD29+) PE-CY7+ (CD56+) cells were sorted out as human MuSCs. Isolated satellite cells were immediately collected or were cultured in Ham’s F10 supplemented with 20% FBS and bFGF (5 ng/ml) (growth medium) for other assays.

### Cells

Mouse C2C12 myoblast cells (CRL-1772) and 293T cells (CRL-3216) were obtained from American Type Culture Collection (ATCC) and cultured in DMEM medium with 10% fetal bovine serum, 100 units/ml of penicillin and 100 μg of streptomycin (growth medium, or GM) at 37 °C in 5% CO2. FACS-sorted MuSCs were cultured in Ham’s F10 medium supplemented with 20% FBS, 5 ng/mL β-FGF (PHG0026, Thermo Fisher Scientific) and 1% P/S at 37°C in 5% CO2 incubator. For PDS treatment, MuSCs or C2C12 cells were treated with DMSO or 5 µM PDS for 48 h then used for EdU incorporation assay, collected for qRT-PCR or library preparation.

### DNA oligonucleotides

All DNA oligonucleotides were purchased from GENEWIZ. The Cy5 fluorescence-labelled DNAs were purified by HPLC. The sequences of the oligos can be found in Supplementary Table S10.

### Plasmids

To study the promoter G4 regions of *Ccne1*, the G4-rich promoter sequences were inserted at the Hind III restriction enzyme site at the upstream of luciferase gene in the plasmid pGL3-Basic (Promega), and the enhancer region was cloned into the BamH I enzyme site at the downstream of firefly. For G4 mutation sequences, all GGG sequences are mutated into GAG and the CCC was replaced by CTC. For AAV-vectored *in vivo* gene editing, the AAV:ITR-U6-sgRNA (backbone)-pCBh-Cre-WPRE-hGHpA-ITR (AAV-Cre) is used as donor plasmid for two sgRNAs insertion. The first selected sgRNA is cloned into the plasmid using Sap I site. The second sgRNA is constructed into the AAV9-single sgRNA vector together with the gRNA scaffold and U6 promoter using Xba I and Kpn I sites. The AAV-dual sgRNA vector is applied in the AAV package. For dCas9-tethered nucleolin system, the sgRNA was inserted into the BsmB I site of plasmid LentiGuide-puro.

### Luciferase reporter assay

Dual luciferase reporter assays were prepared using the Dual-Luciferase Reporter Assay System Kit (Promega, E1910). Typically, 1.5 * 10^5 MuSCs were seeded into each well of a 12-well plate, and after 24h, 500ng wild-type and modified pGL3-Basic plasmids and 20ng pRL were transfected into each well using Lipofectamine 3000. 48h after culturation, the cell medium was discarded and the cells were washed with PBS. 200 μl of lysis buffer was added to the well and incubated at RT for 20 min. Then the cell lysis was centrifuged at 800g for 3 min and the supernatant was used to conduct the luciferase assay according to the manufacturer’s instructions.

### Real-time PCR

Total RNAs from MuSCs or C2C12 cells were extracted using TRIzol reagent (Life Technologies) according to the manufacturer’s protocol. QRT-PCR was performed by using SYBR Green Master Mix (Applied Biosystem) on ABI PRISM 7900HT (Applied Biosystem). *18s rRNA* or *GAPDH* mRNA was used for normalization. All the experiments were designed in triplicates. Primers used for qRT–PCR are shown in Supplementary Table S10.

### CD spectroscopy

CD spectroscopy was performed using a Jasco CD J-150 spectrometer with a 1 cm path length quartz cuvette. The oligos consisting of rG4 sequences were prepared in 10 mM LiCac buffer (pH 7.0) and 150 mM KCl/LiCl. The mixtures, with a reaction volume of 2 mL, were then vortexed and heated at 95°C for 5 minutes and allowed to cool to room temperature. The oligos were examined from 220 to 310 nm at a 2 nm interval, and the data were blanked and normalized to mean residue ellipticity. The data were interpreted using Spectra Manager Suite (Jasco Software).

### Microscale thermophoresis (MST) assay

40nM FAM labelled *Ccne1* G4 or mutated oligos dissolved in the reaction buffer [25mM Tris-HCl (pH=7.5), 150 mM KCl, 1 mM MgCl2] was heated at 75°C for 3 min and slowly cooled down to 4°C. 80 nM of human recombinant MAX protein (Sangon Biotech, C521487) was then added into the sample and incubated at 37°C for 30 min. The samples were loaded to MST capillary tubes, and the blue light mode of the binding software was applied to measure at 25° C. Data were analyzed with Kd mode in the MST nta analysis software.

### EdU assay

The EdU incorporation assay was conducted in accordance with the protocol provided by the Click-iT® Plus EdU Alexa Fluor® 594 Imaging Kit (C10639, Thermo Fisher Scientific). Cells cultured on coverslips were incubated to 10 µM EdU for 4 hours, followed by fixation using 4% paraformaldehyde (PFA) for 15 minutes. The EdU-labeled cells were then detected via click chemistry, employing an Alexa Fluor® 594-conjugated azide. And cell nuclei were stained with DAPI (Life Technologies, P36931). Fluorescent imaging was performed using a fluorescence microscope (Leica).

### AAV packaging

The *in vivo* genome editing via CRISPR/Cas9 was conducted in accordance with previously described instructions. The sgRNA pair targeting MAX was generated from an AAV:ITR-U6-driven and validated *in vitro*. And muscle-tropic AAV9 was employed as the delivery vector.

### Lentivirus packaging and infection

To express dCas9-NCL and sgRNA targeting *Ccne1* G4 sites, lentivirus was generated within 293T cells cultured in the 6-well plate. A mixture of 0.9 μg of pMD2.G, 1.8 μg of psPAX2 and 2.7 μg of pLenti-EF1a-dCas9-nucleolin/pLentiguide-sgRNA/pLentiguide was transfected into each well of 293T cells using 13.5 μl Lipo3000. After 6 h of incubation, the media was replaced with 2 ml DMEM complete medium. Virus particles were harvested at 48 h post-transfection and filtered through a 0.45 μm PES filter, and then stored at -80 °C. For infection of satellite cells, 100 μl virus (50 μl containing dCas9-nucleolin and 50 μl containing sgRNA) were diluted with 400 μl medium with polybrene at a final concentration of 8 μg/ml. After 24 h of infection, the medium was replaced with F10 growth medium, and cells were harvested for experiments after another 24h.

### Immunoblotting and Immunofluorescence

For Western blot assays, *in vitro* cultured cells were harvested, washed with ice-cold PBS and lysed by RIPA lysis buffer with proteinase inhibitor (Thermo Fisher Scientific, 88266) for 15 min on ice. Protein concentration was measured by BCA (Thermo Scientific Pierce BCA Protein Assay Kit, #23227). The protein samples were then loaded to SDS-PAGE. The PVDF membrane with proteins was blocked by 5% BSA for detecting MAX or 5% non-fat milk for other proteins. The following dilutions were used for each antibody: MAX (Cell Signaling Technology, 4739, 1:1000), α-tubulin (Santa Cruz Biotechnology, sc-8654; 1:2000), GAPDH (Sigma-Aldrich, G9545-100UL; 1:4000); and the secondary antibodies: HRP-conjugated Goat anti-Rabbit IgG (ABclonal, AS014, 1:2000), HRP-conjugated Goat anti-Mouse IgG (ABclonal, AS003, 1:2000). Protein expression was visualized using an enhanced chemiluminescence detection system (GE Healthcare, Little Chalfont, UK).

For immunofluorescence staining, cultured cells were fixed in 4% PFA for 15 min and permeabilized with 0.5% NP-40 for 10 min at RT. Then cells were blocked in 3% BSA for 1 h followed by incubating with primary antibodies overnight at 4 °C and secondary antibodies for 1h at RT (protected from light). Then the cells were mounted with DAPI (Life Technologies, P36931) before observation. The appropriate primary antibodies were used as following: PAX7 (Developmental Studies Hybridoma Bank, PAX7-S1ML), MyoD (Dako, M3512), mouse anti-eMyHC (Leica, NCL-MHC-d), and rabbit anti-laminin (Sigma, L9393). For the staining of muscle sections, slides were fixed with 4% PFA for 15 min at room temperature and permeabilized in ice-cold menthol for 6 min at −20 °C. Heat-mediated antigen retrieval with a 0.01 M citric acid (pH 6.0) was performed for 5 min in a microwave. After 4% BBBSA (4% IgG-free BSA in PBS; Jackson, 001-000-162) blocking, the sections were further blocked with unconjugated AffiniPure Fab Fragment (1:100 in PBS; Jackson, 115-007-003) for 30 min. The biotin-conjugated anti-mouse IgG (1:500 in 4% BBBSA, Jackson, 115-065-205) and Cy3-Streptavidin (1:1250 in 4% BBBSA, Jackson, 016-160-084) were used as secondary antibodies. Primary antibodies and dilutions were used as follows PAX7 (Developmental Studies Hybridoma Bank; 1:50), MyoD (Dako; 1:500), eMyHC (Developmental Studies Hybridoma Bank; 1:200) and Laminin (Sigma; 1:800). All fluorescent images were captured with a fluorescence microscope (Leica).

H&E (Hematoxylin-eosin) staining on TA muscle sections was performed according to a protocol described before^14^. Section slides were first stained in hematoxylin for 10 mins followed by rinsing thoroughly under running tap water for at least 3 mins. Then section slides were immersed in 0.2% acid alcohol for 1s and immediately rinsed under running tap water. Next, section slides were stained in eosin for 2 mins followed by rinsing and dehydrating in graded ethanol and Xylene. Finally, slides were mounted by DPX and observed under a normal microscope.

### In cellulo G4 staining

MuSCs were fixed with 4% paraformaldehyde in PBS for 10 min at room temperature. The fixed cells were then permeabilized with 0.5% Triton X-100 in PBS at RT for 20 min followed by incubation with 100 μg/ml RNase A for 1 h at 37 °C and staining with BG4 antibody as previously described^28^.

### PQS prediction

Genomic sequences of mm10 and hg38 were subjected to G4 pattern-matching motifs ^18,52^. A PQS generally consists of at least four G runs (i.e. two or more consecutive Gs) separated by nucleotide stretches of different lengths (loops). The PQS used in this study comply with the following regular expressions:

1. G3+L1–7 = canonical PQS, with at least three tetrads and loops of length up to seven nucleotides: ‘([gG]{3, }\w{1,7}){3, }[gG]{3, }’;
2. G3+L1–12 = extended canonical PQS, with at least three tetrads and longer loops up to 12 nucleotides: ‘([gG]{3, }\w{1,12}){3, }[gG]{3, }’
3. G2L1 – 12 = two-tetrads PQS, with loops up to 12 nucleotides: ‘([gG]{2}\w{1,12}){3,}[gG]{2}’;

### CUT&RUN and data analysis

CUT&RUN assay was conducted using 200,000 satellite cells with the CUT&RUN assay kit (Cell Signaling Technology, 86652,). In brief, FISCs were harvested and washed by cell wash buffer, then bound to concanavalin A-coated magnetic beads. Digitonin Wash Buffer was used for permeabilization. After that, cells were incubated with 2 μg of G4 antibody (Absolute Antibody) overnight at 4 °C with shaking, followed by incubated with 2 μg of anti-flag antibody (Sigma) for 2h at 4 °C with shaking. Then, cell-bead slurry was washed with Digitonin Wash Buffer and incubated with Protein A-MNase for 1 h at 4 °C with shaking. After washing with Digitonin Wash Buffer, CaCl2 was added into the cell-bead slurry to initiate Protein A-MNase digestion, which was then incubated at 4°C for half an hour. Then 2x Stop Buffer was added to the reaction to stop the digestion. CUT&RUN fragments were released by incubation for 30 min at 37 °C followed by centrifugation. After centrifugation, the supernatant was recovered, and DNA purification was performed by using Phenol/Chloroform (Thermo). For DNA library construction, a NEBNext® Ultra™ II DNA Library Prep Kit for Illumina® (NEB, E7645S) was used according to the manufacturer’s instructions. Bioanalyzer analysis and qPCR were used to measure the quality of DNA libraries including the DNA size and purity. Finally, DNA libraries were sequenced on the Illumina Genome Analyzer II platform. The raw data was first pre-processed by initial quality assessment, adapters trimming, and low-quality filtering and then mapped to the mouse reference genome (mm10) or human reference genome (hg38) using Bowtie2^54^, and only the non-redundant reads were kept. The peaks (sites) were identified using MACS2^55^. During the peak calling, the P-value cutoff was set to 0.001 or the Q-value cutoff was set to 0.05 for G4, H3K4me3, and MAX CUT&RUN-Seq experiment. The final result was generated by combining each biological replicate result.

### RNA-seq and data analysis

RNAs were extracted by Trizol reagent and used for library preparation as previously described^27^. Total RNAs were subject to polyA selection (Ambion, 61006) followed by library preparation using NEBNext Ultra II RNA Library Preparation Kit (NEB, E7770S). Libraries were paired-end sequenced with read lengths of 150 bp on Illumina HiSeq X Ten or Nova-seq instruments. The raw reads of RNA-seq were processed following the procedures described in our previous publication. Briefly, the adapter and low-quality sequences were trimmed from 3′ to 5′ ends for each read, and the reads shorter than 50 bp were discarded. The clean reads were aligned to the mouse (mm10) reference genome with Hista2^56^. Next, we used Cufflinks^57^ to quantify the gene expression. Genes were identified as DEGs if the absolute log2foldchange of expression level is greater than 0.5 and the adjusted p-value is <0.05 between two stages/conditions by DESeq2^58^.

### Micro-C

Micro-C for muscle stem cells was performed following the published protocol for mammalian Micro-C^35^. Cells were first crosslinked at 1 ml per million cells of 3mM DSG crosslinker (MedChemExpress, HY-114697) for 35 min at room temperature and then 1% formaldehyde was added and incubated for 10 more minutes. The crosslink was quenched with 0.375 M Tris(pH 7.0) for 5 min. Cells were pelleted at 1000g at 4 °C for 5 min and resuspended with ice-cold PBS. 1 Million cells were split into each tube and pelleted by centrifugation. After snap frozen in liquid nitrogen, cells were kept at -80°C. For cell lysis, cells were thawed on ice for 5 min, and lysed with 0.5 ml MB 1 buffer (10 mM Tris–HCl, pH 7.5, 50 mM NaCl, 5 mM, MgCl2, 3 mM CaCl2, 0.2% NP-40, 1×PIC) on ice for 20 min, washed once with MB 1 buffer, then resuspended in 100 ul MB1 buffer. Mnase concentration for muscle stem cells was predetermined using MNase titration experiments. Chromatin was digested by adding 0.1 ul Mnase (NEB, M0247S) and incubating for 20 min at 37 °C, 1000rpm. Digestion was stopped by adding 8 μl of 500 mM EGTA and incubating at 65 °C for 10 min. Cells were then washed twice with ice-cold MB2 buffer (10 mM Tris–HCl, pH 7.5, 50 mM NaCl, 10 mM, MgCl2). The Mnase digested DNA ends were polished with T4 PNK (NEB M0201), followed by DNA polymerase Klenow fragment (NEB M0210), and then repaired and labeled by adding biotinylated dATP and dCTP (Jena Bioscience, NU-835-BIO14-S and NU-809-BIOX-S, respectively), and TTP/GTP. After washing with buffer MB3 (50 mM Tris–HCl, 10 mM, MgCl2). Ligation was performed for 4 h at room temperature using T4 DNA ligase (NEB M0202). Dangling ends were removed by a 15-minute incubation with Exonuclease III (NEB 0206) at 37 °C. DNAs were de-crosslinked overnight at 65°C. After DNA extraction by ethanol precipitation, size selection was performed using DNA purification beads to enrich ligated fragments at about 230bp. Then biotin selection was done using 10 μl Dynabeads MyOne Streptavidin C1 beads (Invitrogen 65001). Libraries were prepared with the NEBNext Ultra II Library. Preparation Kit (NEB E7103). Samples were paired-end sequenced (read length = 150 bp) on Illumina’s NovaSeq sequencer.

### Micro-C and Hi-C data analysis

The Micro-C raw data was mapped to the mouse reference genome (mm10) using BWA-MEM^59^. Next, we used the parse module of the pairtools^53^ pipeline to find ligation junctions in Micro-C libraries. Then the parsed pairs were then sorted using pairtools sort, and PCR duplication pairs were removed by pairtools dedup. Finally, pairtools split was used to generate .pairs files and .bam files. Loops were identified by HiCCUPS^60^ using parameters (-k KR -t 20) and scaled to 5 kb resolution. Cool files were generated by cooler to further compare the interaction difference. Interaction comparisons were calculated by HiCcompare^61^, ‘A’ value cutoff was set to 15 to filter out interactions with low average expression. Adjusted p value cutoff was set to 0.1 to identify the significant interaction change. The Hi-C raw data was processed by HiC-Pro^62^, and the mouse reference genome was set to mm10. Loops were identified by using parameters (-k KR -t 20) in 5 kb resolution and annotated with histone chip-seq peaks.

### 4C-seq and data analysis

4C-seq was performed as previously described^41^. Approximately 3 million C2C12 cells were cross-linked with 2% (v/v) formaldehyde for 10 min at room temperature and quenched by 0.125 M glycine. The samples were lysed in ice-cold lysis buffer [50 mM tris-HCl (pH=7.5), 150 mM NaCl, 5 mM EDTA, 0.5% NP40, 1% Triton X-100 with complete proteinase inhibitors] on ice for 20 min. Samples were then centrifuged at 700g for 5 min. Pelleted nuclei were washed once with 400 μl 1.2x Dpn II buffer [dilute 10x Dpn II buffer by H_2_O]. The supernatant was discarded, the nuclei were resuspended in 500 μl Dpn II buffer, and 15 μl 10% SDS was added followed by incubation at 37°C for 1 hour with 900 rpm shaking. After incubation. 75 μl of 20% Triton X-100 was added to quench the SDS and then incubated at 37°C for 1 hour with 900 rpm shaking. 200 units of Dpn II restriction enzyme (NEB, R0543) was added, and chromatins were digested at 37°C for overnight. Samples were incubated at 65°C for 20 min to inactivate Dpn II and then cooled to room temperature. 5.7 mL H_2_O, 700 μl 10x NEB T4 DNA ligase buffer and 3350 units of NEB T4 ligase (NEB, M0202) were added to samples and mixed by swirling at room temperature for overnight. 300 μg of proteinase kinase (Invitrogen, AM2548) was added to samples and incubate at 65°C water bath for overnight. 300 μg of RNase A (Thermo Scientific, EN0531) was then added to samples and the samples were incubated at 37°C for 45 min. The samples were subjected to phenol/chloroform/isoamyl extraction and ethanol precipitation and finally resuspended in 150 μl of 10 mM tris-HCl (pH=7.5). The purified DNA samples was then mixed with 50 μl rCutSmart (NEB, B6004), 295 μl H_2_O and 50 units of Nla III (NEB, R0125). The mixture was incubated at 37°C with shaking at 500 rpm shaking on the ThermoMixer overnight. The Nla III restriction enzyme was inactivated by incubating at 65°C for 25 min. A ligation mix containing 12.1 mL H_2_O, 1.4 mL NEB T4 DNA ligase buffer and 100 units of T4 ligase was added to samples, and samples were incubated at room temperature overnight. The samples were purified using phenol/chloroform/isoamyl, precipitated with ethanol, and dissolved in 250 μ l tris-HCl (pH=7.5). Finally, the DNA was then purified with QIAquick PCR Purification Kit (QIAGEN, 28104). A total of 800 ng of purified DNA were used as template for 20 cycles of PCR with Phanta Master Mix (Vazyme, P51101), and the PCR products were purified with NucleoSpin Gel and PCR Clean-up kit (MACHEREY-NAGEL, 740609). The 4C libraries were prepared by on-beads reactions using the NEBNext Ultra II DNA Library Preparation Kit (NEB, E7645S), and sequenced on an NovaSeq X Plus instrument. 150 bp paired-end reads were merged and aligned to mouse genome (mm10) and analyzed with pipe4C^41^. Bigwig file was generated by pipe4C, peak calling was performed with PeakC^63^.

### Statistical test

The statistical significance was assessed by the Student’s two-tailed paired and unpaired t-test. *P<0.05, **P<0.01 and ***P<0.001.

## Author Contributions

X.C. and H.W. conceived the research and designed all the experiments. F.Y. performed all the computational analyses. X.C. and S.Z. performed the CUT&RUN-seq. S.Z. performed mice muscle regeneration analysis upon PDS treatment and MAX KD. X.G. conducted the luciferase reporter assay, AAV package and injection, CRISPR-dCas9 tethering assay. J.Z. conducted the CD assay and MST assay. Y.Q. conducted Micro-C. L.H. supervised the AAV package and injection. Y.L. assisted in the collection of human muscle tissues. Q.Z. assisted in the FACS sorting of muscle stem cells. M.O. collected human muscle tissues. H.S. supervised computational analyses. C.K.K. supervised the validation of G4 structure and interaction between G4 and MAX. X.C., F.Y. and H.W. wrote the manuscript with input from all authors.

## Acknowledgements

This work was supported by the National Natural Science Foundation of China (NSFC) [32300703 to X.C.]; Natural Science Foundation of Guangdong Province, China [2024A1515030291] to X.C.; National Key R&D Program of China [2022YFA0806003 to H.W.]; National Natural Science Foundation of China [82172436 to H.W.]; General Research Fund (GRF) from the Research Grants Council (RGC) of the Hong Kong Special Administrative Region, China [14106521, 14100620, 14105823 and 14115319 to H.W.;14103522, 14105123 and 14120420 to H.S.]; Theme-based Research Scheme (TRS) from RGC [T13-602/21-N to H.W.]; Strategic Topics Grant (STG) from RGC [STG1/E-403/24-N to H.W.]; Area of Excellence Scheme (AoE) from RGC [AoE/M-402/20 to H.W.]; Health and Medical Research Fund (HMRF) from Health Bureau of the Hong Kong Special Administrative Region, China [10210906 and 08190626 to H.W.]; the research funds from Health@InnoHK program launched by Innovation Technology Commission, the Government of the Hong Kong SAR, China to H.W.; Chinese University of Hong Kong (CUHK) Strategic Seed Funding for Collaborative Research Scheme (SSFCRS) to H.W..

---

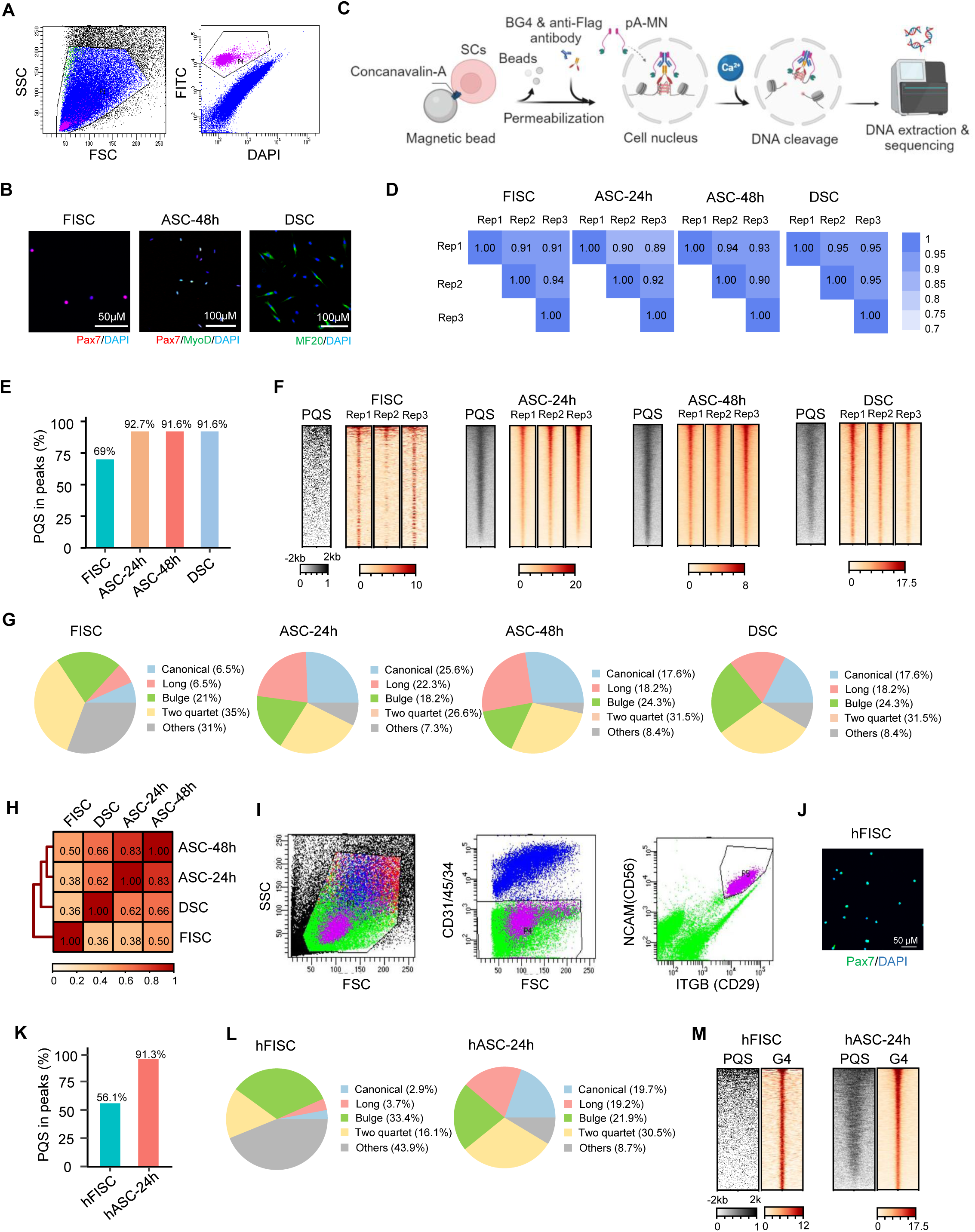

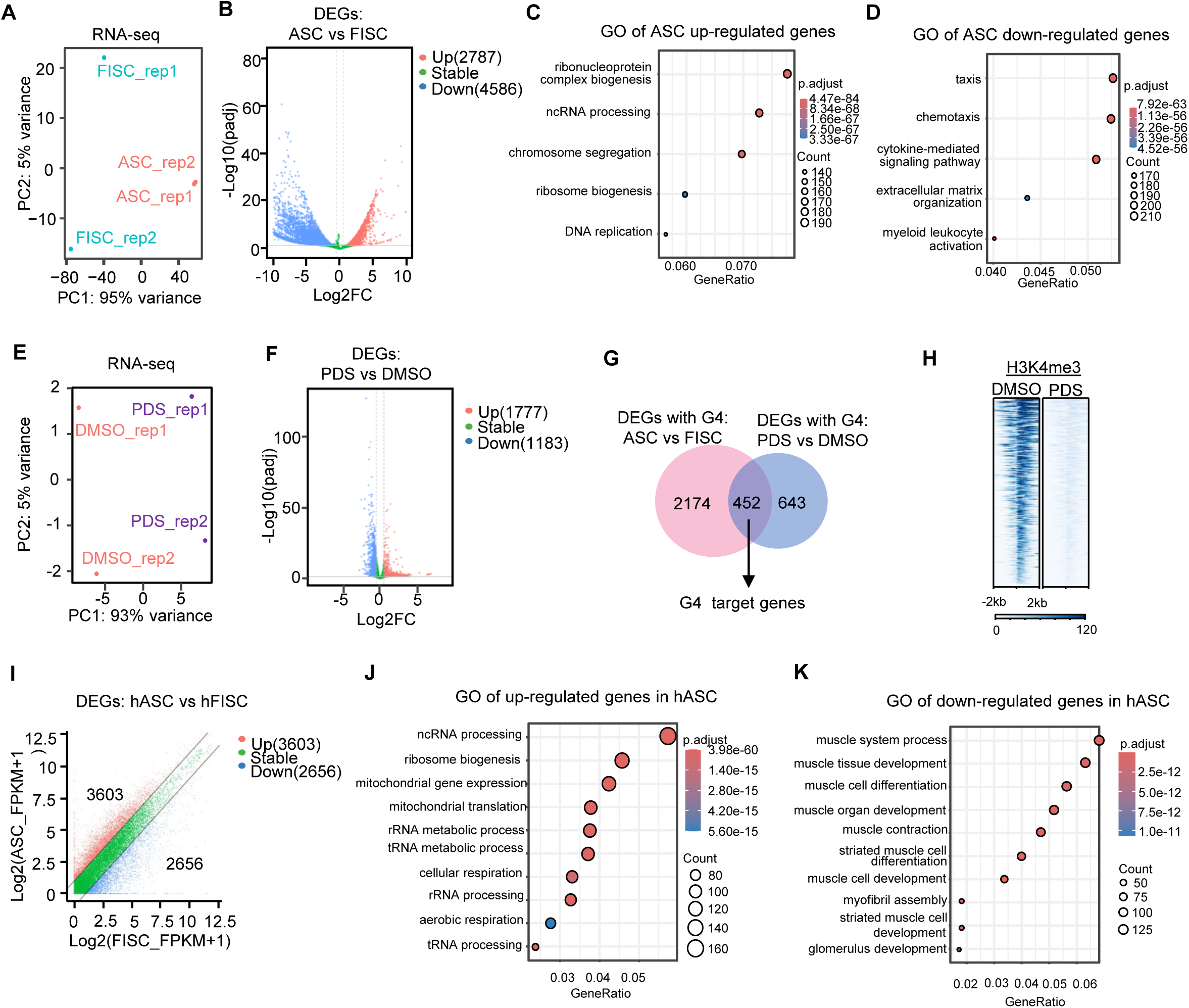

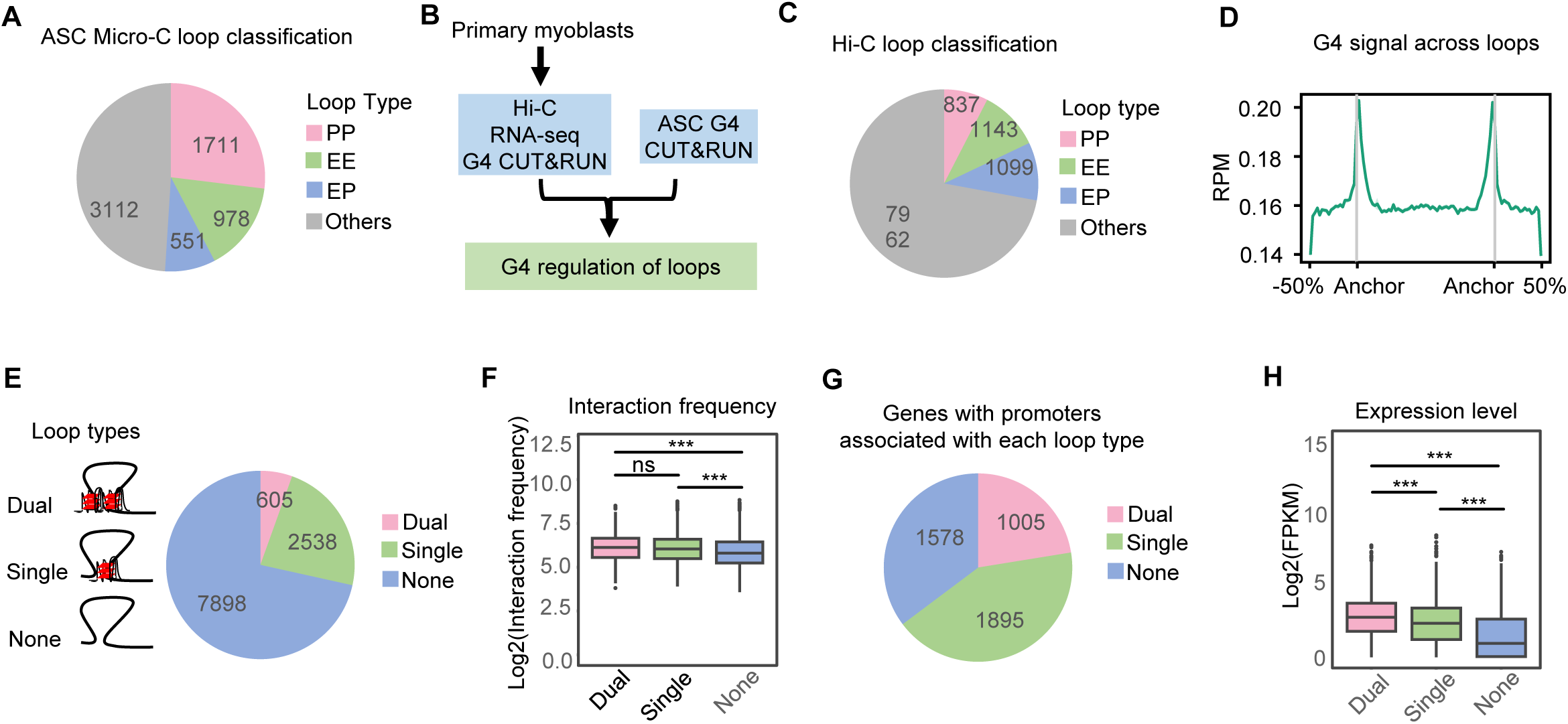

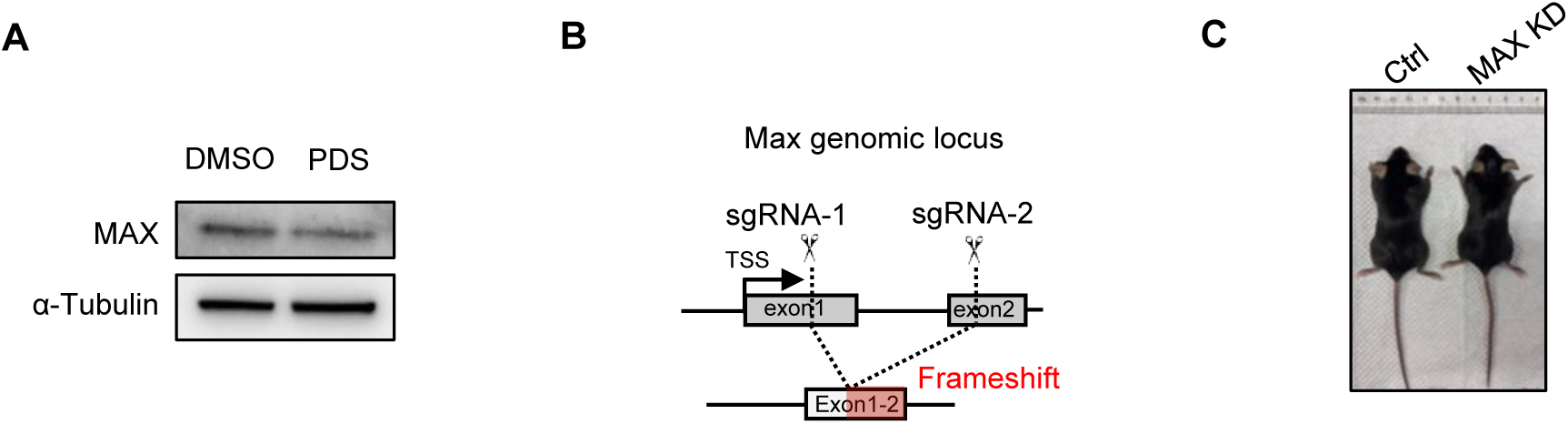

