## Supplemental Table S1-S10 for "DNA G-quadruplex Profiling Reveals Functional and Mechanistic Role of G-quadruplexes in Skeletal Muscle Stem Cells": Suppl_Info.docx

**Inventory of Supplementary Information**

1. **Supplementary Figures**

Suppl. Fig. S1. G4 profiling reveals dynamic remodeling of G4s during MuSC lineage progression.

Suppl. Fig. S2. Promoter G4 formation regulates gene transcription in ASCs.

Suppl. Fig. S3. G4s are enriched at loop anchors and promote loop interactions in ASCs.

Suppl. Fig. S4. MAX promotes MuSCs proliferation and adult muscle regeneration.

2. **Supplementary Tables**

Suppl. Table S1. G4 CUT&RUN profiling in MuSCs.

Suppl. Table S2. G4-containing promoters in MuSCs.

Suppl. Table S3. RNA-seq analysis in FISCs, ASCs and DMSO or PDS treated ASCs.

Suppl. Table S4. H3K4me3 CUT&RUN analysis in ASCs treated with DMSO or PDS.

Suppl. Table S5. RNA-seq analysis in human FISCs and ASCs.

Suppl. Table S6. Micro-C analysis in ASCs treated with DSMO or PDS.

Suppl. Table S7. ChIP-seq analysis of TFs in C2C12 cells.

Suppl. Table S8. MAX CUT&RUN analysis in ASCs.

Suppl. Table S9. Micro-C and RNA-seq analysis in ASCs transfected with siNC or siMAX oligos.

Suppl. Table S10. Information of oligonucleotides and primers used in the study.

**3. Supplementary Figure Legends**

**Suppl. Fig. S1. G4 profiling reveals dynamic remodeling of G4s during MuSC lineage progression.** (A) Mouse MuSCs were isolated from PAX7-nGFP mice and FACS gating images are shown. (B) Schematic illustration of G4 CUT&RUN-seq procedure. (C) IF staining of PAX7 (red) in FISCs, PAX7 (red) and MyoD (green) in ASC-48h and MF20 (green) in DSCs to validate the cell identify at each stage during the lineage progression. Nuclei were visualized by DAPI staining (blue). (D) G4 CUT&RUN was performed with three replicates from each of the above stages. Heatmaps showing signal correlation of shared G4 peaks from the three replicates. Pearson correlation coefficient was calculated. (E) The Boxplot showing the fraction of PQS from the above identified G4 peaks. (F) Heatmaps showing the PQS and G4 signal enrichment in the identified G4 peaks. (G) Pie chart showing the distribution of G4 subtypes in the above identified G4s. (H) Heatmap showing the correlation of G4 signals in the four stages. Pearson correlation coefficient was calculated. (I) Human MuSCs were isolated by FACS with staining of antibodies (CD31-/CD45-/CD34-/ITGB1+/NCAM+) and the gating images are shown. (J) IF staining of Pax7 (green) to validate the above isolated hFISC cell identity. Nuclei were visualized by DAPI staining (blue). (K) Boxplot showing the fraction of PQS in the identified G4 peaks in hFISC and hACS-24h. (L) Distribution of G4 subtypes in the above identified G4 peaks in hMuSCs. (M) Heatmaps showing the PQS and G4 signal enrichment in the above identified G4 peaks in hMuSCs.

**Figure S2. Promoter G4 formation regulates gene transcription in ASCs**. (A) RNA-seq was performed in mouse FISCs and ASCs. Principal component analysis (PCA) of the variance-stabilized estimated raw counts of differentially expressed genes. Each dot represents an individual biological replicate. (B) Volcano plot showing differentially expressed genes identified in mouse ASC vs. FISC. The red and blue dots denote up- and down-regulated genes, and the green dots denote unchanged genes. (C-D) GO analysis of the above identified up- and down-regulated genes in ASCs. (E) ASCs were treated with PDS or DMSO (control) and RNA-seq was performed. PCA of the variance-stabilized estimated raw counts of differentially expressed genes. Each dot represents an individual biological replicate. (F) Volcano plot showing differentially expressed genes identified in ASCs treated with PDS vs. DMSO. (G) Comparison of DEGs from ASC vs. FISCs and DEGs from ASCs treated with PDS vs. DMSO to identify potential G4 target genes. (H) H3K4me3 signals at the promoter of G4 activated genes in ASCs treated with DMSO or PDS. (I) RNA-seq was performed in hFISCs and hASCs. Scatter plot showing differentially expressed genes in hASCs vs. hFISCs. (J-K) GO analysis of the above identified up- and down-regulated genes.

**Figure S3. G4s are enriched at loop anchors and promote loop interactions in ASCs.** (A) Micro-C was performed in ASCs and chromatin loops were identified. Pie chart showing the number of each type of loops. PP: promoter-promoter loop; EE: enhancer-enhancer loop; EP: enhancer-promoter loop; others: none of the above. (B) Schematic illustration of integrative analysis to investigate G4 regulation of chromatin looping in primary myoblast cells integrating the publicly available RNA-seq, Hi-C and H3K27ac ChIP-seq datasets. (C) Loops of different subtypes identified in the above Hi-C data at 5k resolution. (D) G4 signals across the above identified loops. (E) Classification of the above identified loops based on G4 localization at the loop anchors. Pie chart showing the number of each type of G4-related loops. (F) Comparison of interaction frequency of the above different type of G4 loops. Two-tailed Student’s t-test was used for statistical calculation: ns P>0.05, ***P < 0.001. (G) The number of genes with promoters associated with each type of G4 loops. (H) Expression level of genes with promoters associated with each type of G4 loops. Two-tailed Student’s t-test was used for statistical calculation: ***P < 0.001.

**Figure S4. MAX promotes MuSCs proliferation and adult muscle regeneration.** (A) MAX protein level detected by western blot in ASCs treated with DMSO or PDS. α-Tubulin was used as the normalization control. (B) Schematic of the strategy for MAX inactivation using CRISPR-Cas9 editing in Pax7^Cas9^ mice. Two sgRNAs were designed to target the exon 1 and exon 2 of MAX ORF, respectively to achieve frameshift mutation of MAX in MuSCs. (C) No obvious morphological difference was observed in Ctrl vs. MAX KD mice.
